# Rapid Identification of Neutralizing Antibodies against SARS-CoV-2 Variants by mRNA Display

**DOI:** 10.1101/2021.09.14.460356

**Authors:** Shiho Tanaka, C. Anders Olson, Christopher O. Barnes, Wendy Higashide, Marcos Gonzalez, Justin Taft, Ashley Richardson, Marta Martin-Fernandez, Dusan Bogunovic, Priyanthi N.P. Gnanapragasam, Pamela J. Bjorkman, Patricia Spilman, Kayvan Niazi, Shahrooz Rabizadeh, Patrick Soon-Shiong

## Abstract

The increasing prevalence of SARS-CoV-2 variants with the ability to escape existing humoral protection conferred by previous infection and/or immunization necessitates the discovery of broadly-reactive neutralizing antibodies (nAbs). Utilizing mRNA display, we identified a set of antibodies against SARS-CoV-2 spike (S) proteins and characterized the structures of nAbs that recognized epitopes in the S1 subunit of the S glycoprotein. These structural studies revealed distinct binding modes for several antibodies, including targeting of rare cryptic epitopes in the receptor-binding domain (RBD) of S that interacts with angiotensin-converting enzyme 2 (ACE2) to initiate infection, as well as the S1 subdomain 1. A potent ACE2-blocking nAb was further engineered to sustain binding to S RBD with the E484K and L452R substitutions found in multiple SARS-CoV-2 variants. We demonstrate that mRNA display is a promising approach for the rapid identification of nAbs that can be used in combination to combat emerging SARS-CoV-2 variants.

## Introduction

The emergence of severe acute respiratory syndrome coronavirus 2 (SARS-CoV-2), the causative agent of the respiratory disease COVID-19, has resulted in a pandemic that brought the world to a standstill (Zhou et al., 2020). Despite the rapid development and success of vaccines and antibody therapies, ongoing SARS-CoV-2 antigenic drift has resulted in the emergence of variants that pose new threats (Davies et al., 2021; Plante et al., 2021; Yurkovetskiy et al., 2020). Various studies have shown that several of these variants have the ability to escape antibody neutralization mediated by antisera from recovered COVID-19 patients/vaccinated individuals or recombinant neutralizing antibodies (nAbs) developed as therapeutics (Cerutti et al., 2021; McCallum et al., 2021a; Suryadevara et al., 2021). Thus, along with modified vaccines to combat variants, there is an urgent need for development of prophylactic and therapeutic anti-viral drugs, including biologics such as nAbs, with sustained efficacy against SARS-CoV-2 variants.

The trimeric SARS-CoV-2 spike (S) glycoprotein serves as the fusion machinery for viral entry, and therefore represents the main target of nAbs (Brouwer et al., 2020; Cao et al., 2020; Robbiani et al., 2020). The SARS-CoV-2 S trimer utilizes the angiotensin-converting enzyme 2 (ACE2) as its host receptor (Hoffmann et al., 2020; Li et al., 2003; Zhou et al., 2020), through interactions with the receptor-binding domains (RBDs) located at the apex of the S trimer. The RBDs adopt either ‘down’ or ‘up’ conformations, with RBD binding to ACE2 facilitated only by the ‘up’ conformation (Kirchdoerfer et al., 2016; Li et al., 2019; Walls et al., 2016, 2020; Wrapp et al., 2020; Yuan et al., 2017). While the majority of potent anti-SARS-CoV-2 nAbs target the RBD and directly compete with ACE2 binding (Barnes et al., 2020a; Brouwer et al., 2020; Cao et al., 2020; Robbiani et al., 2020), recent studies have revealed nAbs that target the N-terminal domain (NTD) (Liu et al., 2020; McCallum et al., 2021b) and S2 stem helix (Zhou et al., 2021).

The structures of numerous monoclonal antibodies (mAbs) recognizing the RBD and NTD have been characterized (Barnes et al., 2020b, 2020a; Baum et al., 2020; Brouwer et al., 2020; Hansen et al., 2020; Pinto et al., 2020), enabling their classification based on shared epitopes and neutralizing properties (Barnes et al., 2020b; Dejnirattisai et al., 2021; McCallum et al., 2021b; Yuan et al., 2021). A subset of mAbs that recognize non-overlapping epitopes are in clinical trials or have received emergency use authorization from the US Food and Drug Administration (FDA) for the treatment and prevention of COVID-19 (Cathcart et al., 2021; Jones et al., 2021; Weinreich et al., 2021). However, ongoing viral evolution and genetic drift has resulted in an accumulation of mutations and/or deletions found in the S RBD and NTD that enhance affinity of ACE2 binding and allow some variants to evade existing immunity (Cele et al., 2021; Tegally et al., 2021). Thus, current emergency-authorized therapies developed early in the pandemic based on the first-wave or ‘A’ strain S sequence could potentially be less effective against emerging SARS-CoV-2 variants that harbor escape mutations mapped to their epitopes (Greaney et al., 2021a, 2021b; Starr et al., 2020, 2021; Weisblum et al., 2020).

Here, we report our identification via mRNA display (Newton et al., 2020; Olson et al., 2008; Roberts and Szostak, 1997; Takahashi et al., 2003) of a set of novel mAbs targeting SARS-CoV-2 S, which we demonstrate neutralize both authentic and pseudoviral SARS-CoV-2 with IC50s between 0.076 – 7.0 µg/mL. Structural analysis revealed a subset of these nAbs recognize RBD and NTD epitopes, including a rare, cryptic, cross-reactive RBD epitope.

Moreover, we characterize a weakly neutralizing antibody that recognizes the S1 subdomain 1 (SD1), providing insight into a unique class of antibodies that are infrequently found among convalescent individuals (Zost et al., 2020a, 2020b) that can be utilized in the fight against COVID-19. Finally, we describe the utility of mRNA display for rapid identification of variant-resistant antibody clones. This powerful technique enabled the rapid selection of a discovered SARS-CoV-2 nAb to extend its neutralizing capability to SARS-CoV-2 expressing the E484K and L452R S RBD mutations found in multiple SARS-CoV-2 variants.

## Results

### Identification of Anti-SARS-CoV-2 Spike Antibodies by mRNA Display

We utilized mRNA display to identify mAbs targeting the S protein of SARS-CoV-2 and discovered 10 novel VH/VL sequences that bind various domains on S (Table S1). S comprises an N-terminal fragment known as S1, which further divides into the NTD, ACE2 RBD, small C- terminal subdomains 1 and 2 (SD1 and SD2), and a C-terminal “S2” fragment (Figure 1A). Bio- layer interferometry (BLI) kinetic analysis using recombinant SARS-CoV-2 RBD (residue 319- 541), RBD-SD1 (residue 319-591), S1 (residue 16-685), and S2 (residue 686-1213) proteins revealed 3 antibodies bind the RBD (N-612-017, N-612-056, and N-612-074), 2 antibodies bind the SD1 domain (N-612-004 and N-612-041), 2 antibodies bind the NTD (N-612-002 and N- 612-014), and 3 antibodies bind the S2 domain (N-612-007, N-612-044, and N-612-086) (Figure 1A-C). All 10 antibodies bind corresponding binding domains with low nM binding affinity (*K*D) (Figure 1B and Table S2), but apparent affinities are far superior (*K*D < 3 pM) for S trimers (Figure 1B and 1C, and Table S3). Among the 3 RBD binders, only N-612-017 showed competition with ACE2 binding (Figure 1D).

**Figure 1.**
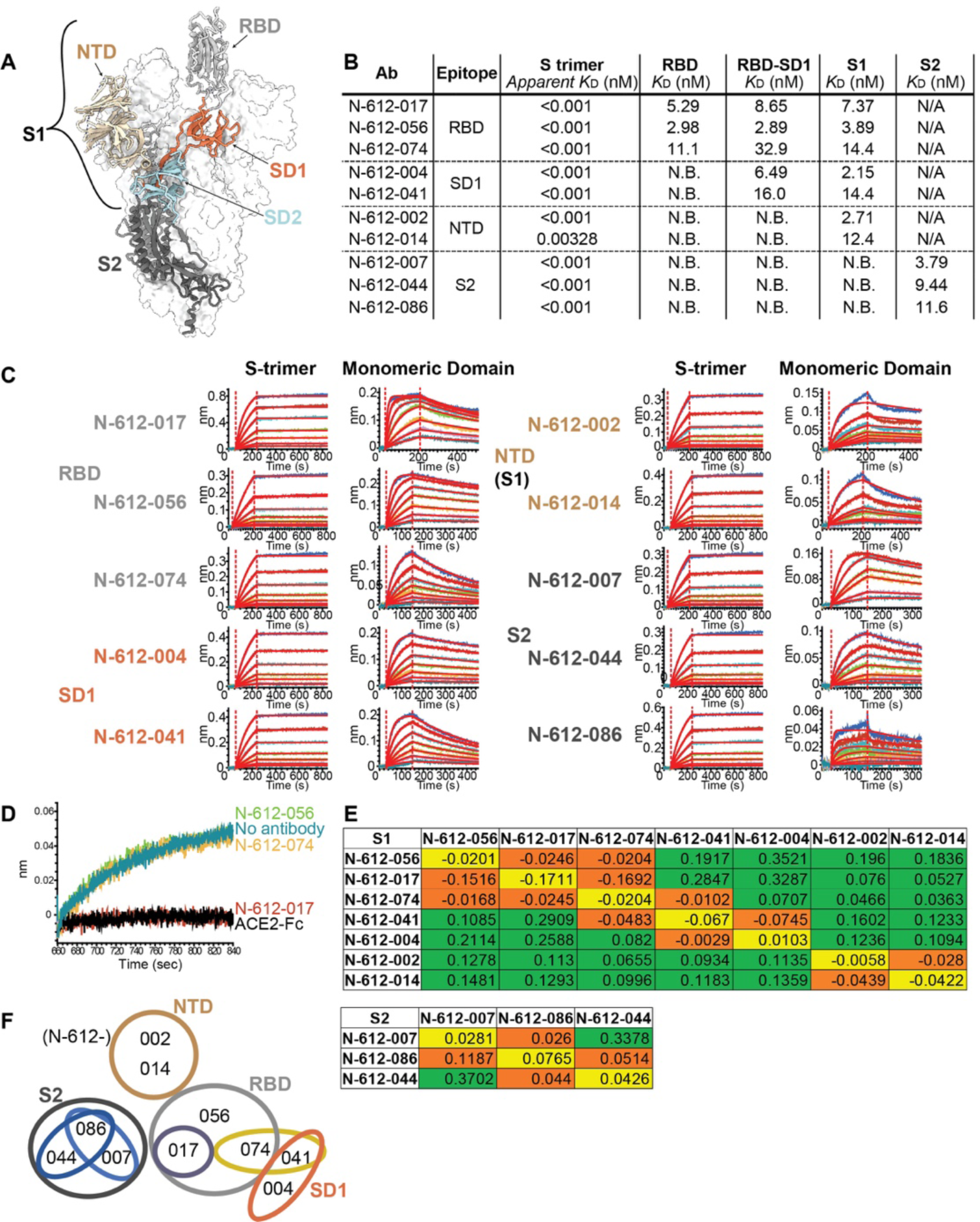
Identification of SARS-CoV-2 Spike Targeting Monoclonal Antibodies. (A) Model of the SARS-CoV-2 spike trimer domains (PDB 6VYB); NTD (wheat), RBD (light gray), SD1 (coral), SD2 (powder blue), and S2 (dark blue). (B) *K*D summary table from BLI kinetic analysis of 10 antibodies against spike trimer and various domains used as analytes. N.B. indicates no binding. N/A is untested. Apparent *K*D values for S-trimer were obtained by curve fitting with a bivalent model. (C) BLI kinetic analysis of 10 antibodies against the spike trimer (left) and each corresponding domain (right). (D) BLI blocking assay: biosensors were coated with RBD and subsequently all RBD binding antibodies (N-612-017, N-612-056, and N-612-074) and ACE2- IgG1Fc were incubated with RBD coated biosensor. The recorded signal from ACE2-IgG1Fc binding to the RBD on the biosensor indicates the RBD blocking capability of the test samples. Both N-612-017 and ACE2-IgG1Fc completely blocked RBD and ACE2 interaction. (E) Epitope binning data indicating competing antibody pairs in red and non-competing antibody pairs in green. Self-blocking is in orange. (F) Epitope binning diagram mapping overlapping regions of binding sites of 10 mAbs.

To further map the binding regions of the 10 antibodies, we performed epitope binning experiments using S1 and S2 fragments separately (Figure 1E). Two NTD binders blocked each other but not RBD or SD1 binding antibodies. The 3 RBD binders competed with each other (Figure 1E and F) despite N-612-017 being the only ACE2 blocker (Figure 1D). SD1 binders N- 612-004 and N-612-041 blocked each other, but only N-612-041 blocked N-612-074 (an RBD binder) suggesting N-612-041 and N-612-074 have proximal or overlapping binding sites (Figure 1E and 1F). Epitope binning using the S2 domain revealed N-612-007 and N-612-044 are non-competing whereas N-612-086 competes with both N-612-007 and N-612-044, suggesting they all bind distinct epitopes on S2 (Figure 1E and 1F).

In addition, multiple biophysical assays were carried out to determine the developability of all 10 antibodies (Table S4) (Jain et al., 2017). All 10 mAbs displayed low polyreactivity scores by meso scale diagnostic (MSD) analysis and low self-interaction scores by BLI-clone self-interaction (CSI) (Table S4). Eight of the mAb candidates exhibited low hydrophobicity in the hydrophobic interaction column (HIC) chromatography while higher hydrophobicity was observed for N-612-041 and N-612-074 (Table S4). N-612-041 also showed more rapid aggregation in an accelerated stability assay system while the other 9 mAbs demonstrated long-term stability (Table S4). Furthermore, all 10 mAbs exhibited desirable thermostability of Fab in differential scanning fluorimetry (DSF) melting temperature (Tm) analysis, although N-612-044 exhibited heterogeneous characteristics in thermostability and hydrophobic interaction column chromatography. Ultimately, 7 out of 10 antibodies displayed biophysical characteristics within the acceptance criteria, indicating antibodies engineered by mRNA display can have favorable developability.

### Neutralization Activity Assessment of Anti-SARS-CoV-2 Antibodies

Ten mAbs identified by mRNA display were assessed for neutralization activity against authentic SARS-CoV-2 virus in a Vero E6 cell neutralization assay. The ACE2-blocking anti-RBD antibody N-612-017 demonstrated the highest neutralization activity and the non-ACE2- blocking RBD binder N-612-056 showed weaker but nearly complete neutralization of ∼87%. (Figure 2A). N-612-004 (SD1 binder), N-612-007 (S2 binder), and N-612-014 (NTD) antibodies all showed some neutralization activity that plateaued at 40∼60% (Figure 2A), similar to previous observations made for anti-NTD antibodies (McCallum et al., 2021b). We next investigated the activity of N-612-017 in combination with N-612-004, N-612-007, and N-612-014. N-612-017 and respective partners were mixed in equal concentrations. All combinations tested showed slightly improved IC50 values compared to N-612-017 by itself, suggesting both antibodies present in mixture can bind to S simultaneously and in some cases non-RBD domain binders can enhance activity of RBD-binding nAb (Figure 2B, C and D).

**Figure 2.**
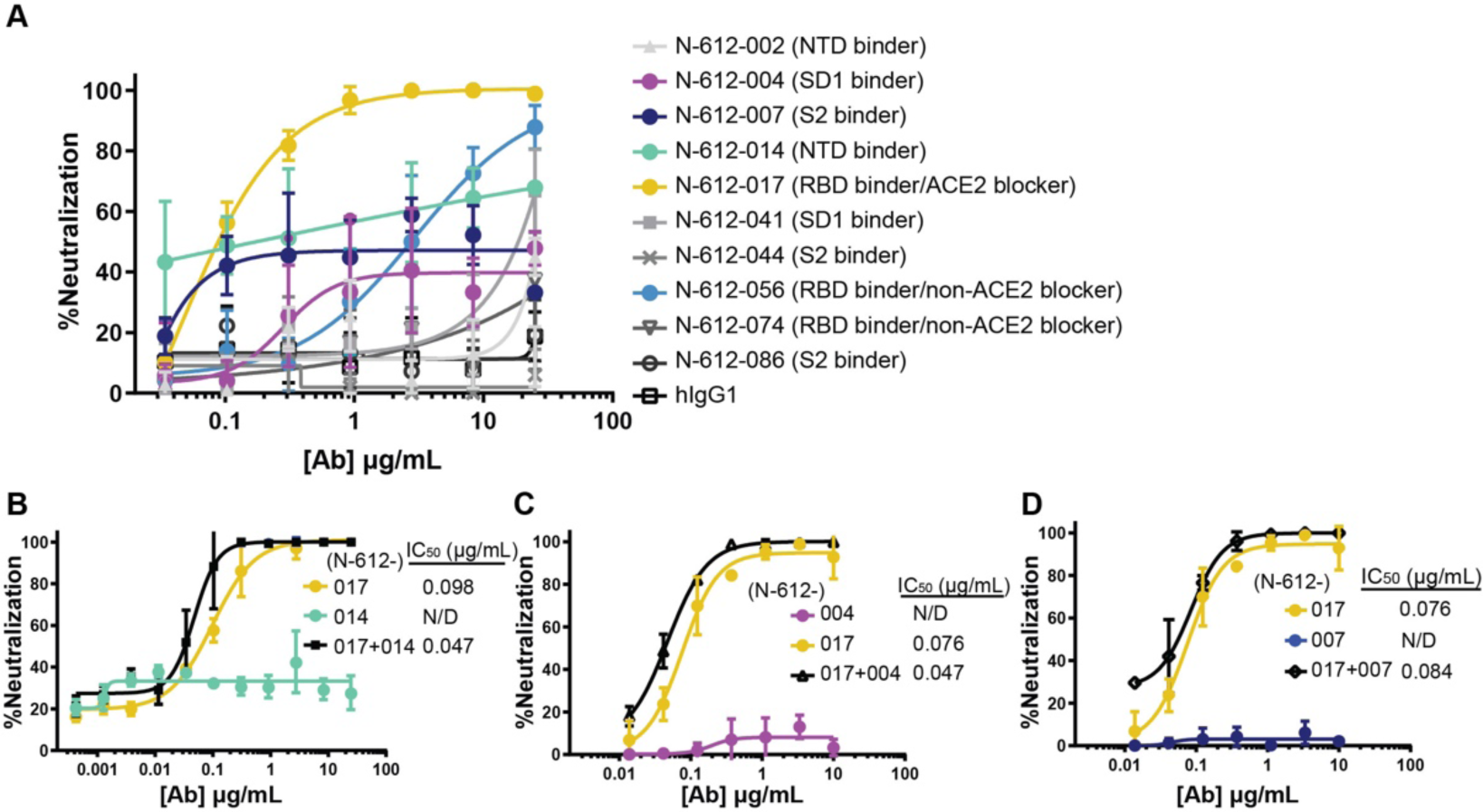
Neutralization Activity of mAbs in Vero E6 Live Virus Neutralization Assay. (A) Dose-dependent neutralization of SARS-CoV-2 virus by 10 mAbs selected by mRNA library display. N-612-017 neutralization activity in combination with (B) N-612-014, (C) N-612-004, and (D) N-612-007. X-axis represents the concentration of antibodies when used alone; when antibodies were combined, an equal concentration of each antibody was used.

Neutralization activity of N-612-014 (NTD binder) showed variable saturation between assays and prevented accurate IC50 determination (Figure S1A). To test whether activity of N- 612-014 changes in time-dependent manner, we tested the effects of longer antibody-virus incubation times on neutralization potency. With a virus-antibody incubation time of ∼30 min, neutralization activity plateaued between 20∼90% (Figure S1A). In contrast, when virus was incubated with antibody for 24 hours, neutralization activity plateaued at ∼90% with IC50 values of 0.023-0.025 µg/mL (Figure S1B). Longer incubation also resulted in improved neutralization potencies for N-612-017, N-612-056, and positive control convalescent plasma serum (Figure S1C), suggesting a change in viral infectivity due to a time-dependent conformational change in spike (Huo et al., 2020; Wec et al., 2020).

### Structural Characterization of RBD-Specific, ACE2 Blocking nAb N-612-017

To investigate the specificity of RBD-targeting for nAbs N-612-017 and N-612-056, we determined a 3.2 Å single-particle cryo-electron microscopy (cryo-EM) structure of a complex between SARS-CoV-2 S trimer and the N-612-017 Fab (Figure 3, Figure S2 and Table S5), and a 2.9 Å X-ray crystal structure of a SARS-CoV-2 RBD – N-612-056 Fab complex (Figure 4 and Table S6). The N-612-017 – S trimer complex structure revealed N-612-017 Fab binding to both ‘up’ and ‘down’ RBD conformations, and recognition of an epitope that partially overlapped with the ACE2 receptor binding site (Figure 3A and 3B), consistent with BLI competition data (Figure 1D). N-612-017 uses five of its six complementarity-determining region (CDR) loops and HC framework region 3 (FWR3) to interact with an epitope focused on RBD residues adjacent to the ACE2 receptor binding ridge (Figure 3C and 3D), resulting in ∼1018Å^2^ buried surface area (BSA) on the epitope. The CDRH2 and CDRH3 loops mediate the majority of RBD contacts (∼616Å^2^ of ∼1030Å^2^ total paratope BSA), establishing hydrophobic and hydrogen bond interactions at the Fab-RBD interface. Of note, N-612-017 CDRH2 loop residues contact RBD positions frequently mutated among circulating variants (Deng et al., 2021; Kuzmina et al., 2021; McCallum et al., 2021a; Wang et al., 2021). RBD residue E484RBD established hydrogen bond interactions with G52HC and G54HC in CDR2 and D72HC in FWR3 (Figure 3E), while L452RBD formed stacking interactions with CDR2 residue Y52HC (Figure 3F). Taken together, these data indicate nAb N-612-017 targets the RBD similarly to nAbs that belong to the class 2 binding mode, which is the predominant nAb class identified in convalescent and vaccinated donors (Barnes et al., 2020b; Wang et al., 2021).

**Figure 3.**
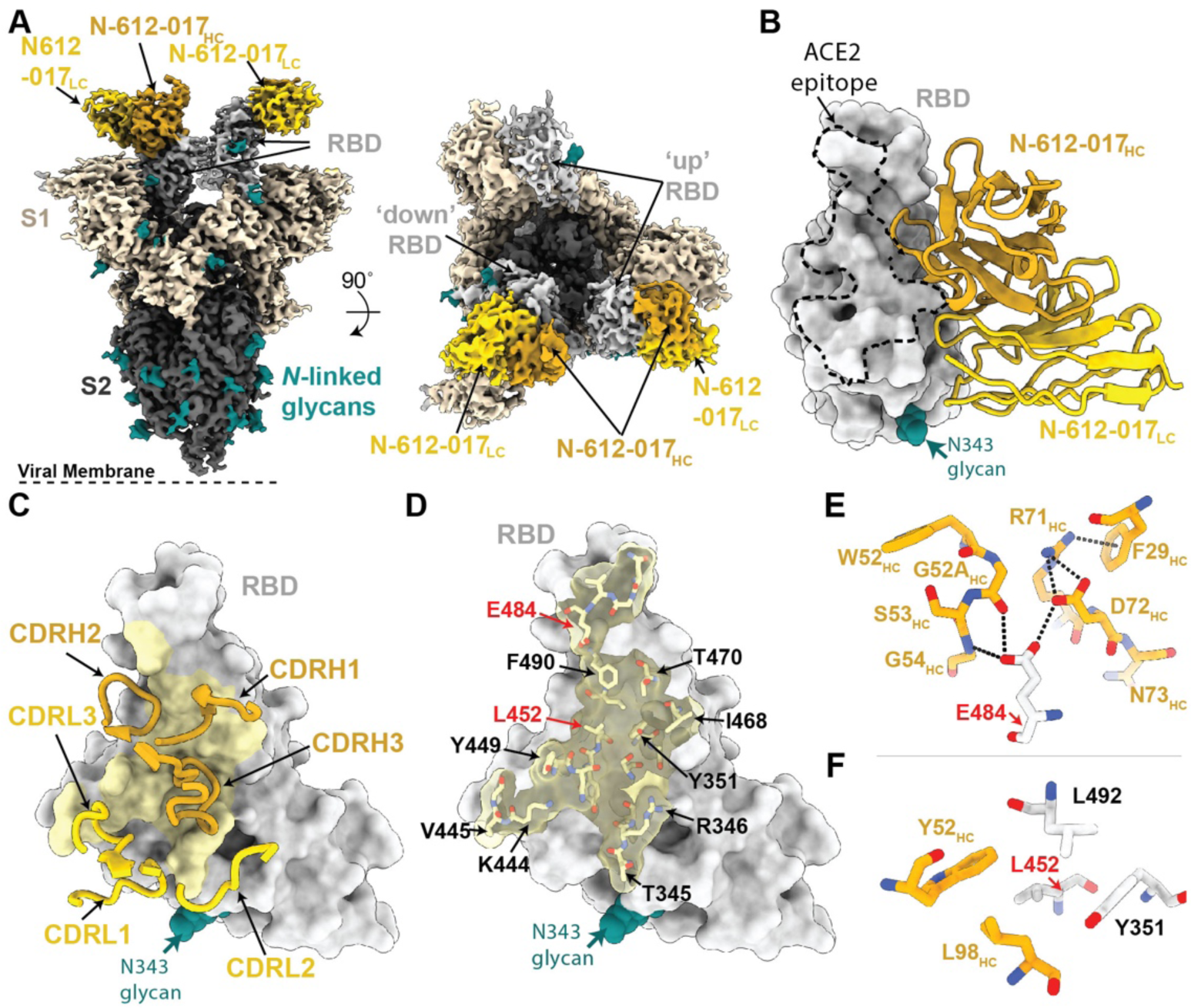
Cryo-EM Structure of the N-612-017 – S Complex. (A) Cryo-EM density for the N- 612-017- S trimer complex. Side view (left panel) illustrates orientation with respect to the viral membrane (dashed line). (B) Close-up view of N-612-017 variable domains (HC: gold, LC: yellow) bound to RBD (gray surface). ACE2 receptor binding site is shown as a dashed line. (C) N-612-017 CDR loops mapped on the RBD. (D) Surface and stick representation of N-612-017 epitope (yellow) on RBD surface (gray). (E,F). Residue-level interactions between N-612-017 (gold) and SARS-CoV-2 RBD (gray). Potential hydrogen bond interactions are illustrated by dashed black lines.

**Figure 4.**
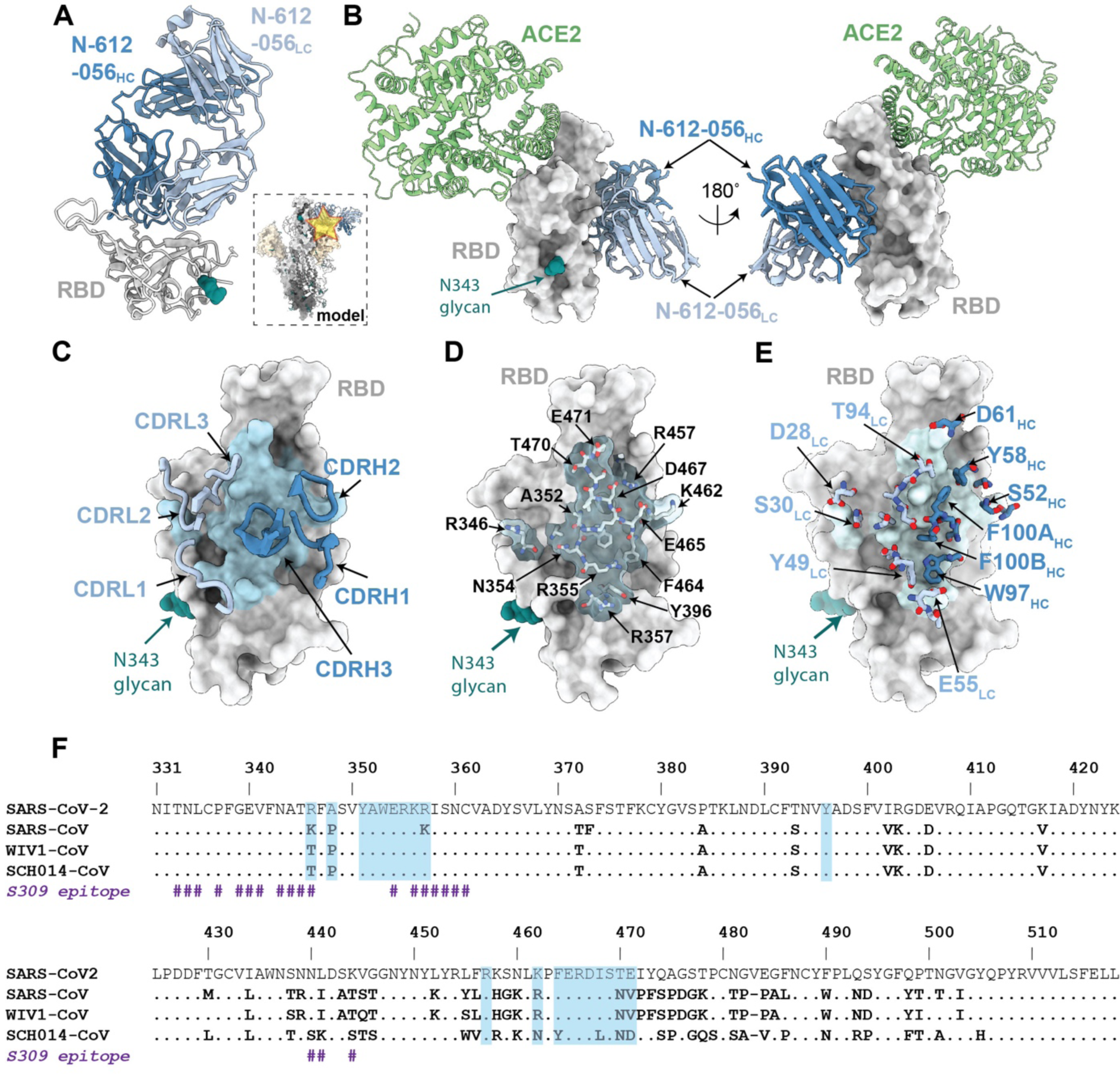
X-ray Crystal Structure of SARS-CoV-2 RBD in Complex with the N-612-056 Fab. (A) 2.9 Å X-ray crystal structure for the N-612-056 Fab – RBD complex. Inset: Overlay of the N-612-056-RBD crystal structure on a S trimer with ‘up’ RBD conformation (PDB 6VYB). (B) Composite model of N-612-056 – RBD (blue ribbon and gray surface, respectively) overlaid with soluble ACE2 (green, PDB 6M0J). The model was generated by aligning RBDs on 191 matched C*α* atoms. (C) N-612-056 CDR loops (blue) mapped on the RBD surface (gray). The N-612-056 epitope is shown as a light blue surface. (D) Surface and stick representation of N- 612-056 epitope. (E) N-612-056 paratope residues mapped on the RBD surface with epitope residues shown in light blue. (F) Sequence alignment of SARS-CoV-2, SARS-CoV, WIV1-CoV, and SCH014-CoV. N-612-056 epitope residues are shaded blue. S309 epitope residues are also shown (# symbol).

### Structural Characterization of RBD-Specific nAb N-612-056 Targeting Cryptic Site

Next, we analyzed the high-resolution X-ray crystal structure of the SARS-CoV-2 RBD –N- 612-056 Fab complex (Figure 4A). This method was used rather than cryo-EM due to N-612- 056’s lack of binding to intact S trimers (Figure 4A; inset). Similar to the donor-derived antibody COVOX-45 (Dejnirattisai et al., 2021), N-612-056 binds a rare cryptic epitope that is not readily found in the repertoire of antibodies from convalescent donors (Figure S4). Consistent with observed binding to dissociated S1 protomers by single-particle cryo-EM (data not shown), the N-612-056 cryptic epitope is inaccessible on an S trimer due to steric clashes with the neighboring NTD, and does not overlap with the ACE2 binding site (Figure 4A and 4B). N-612- 056 HC and LC CDR loops participate equally to bury ∼890 Å^2^ of the RBD epitope surface area that comprises residues 352-357 in the *β*1 strand, which is part of a structurally conserved 5- stranded RBD *β*-sheet, and residues 457-471 that comprise a disordered loop directly beneath the ACE2 receptor binding ridge (Figure 4C and 4D).

N-612-056 establishes a network of hydrogen bond and hydrophobic interactions that include a stretch of hydrophobic residues in CDRH3 that mediate van der Waals interactions at the RBD interface, and the formation of salt bridges between N-612-056 residues D28LC and E55LC with R346RBD and R357RBD, respectively (Figure 4E). These structural data explain the observed cross-reactivity against SARS-CoV RBD (Figure S3A)(Cohen et al., 2021), as 14 of 20 epitope residues are strictly conserved and three additional residues (R346, R357 and K462 SARS-CoV-2 RBD numbering) are conservatively-substituted (K333, K344, and R449 SARS-CoV RBD numbering) in SARS-CoV and SARS-CoV-2. (Figure 4F). Overall, these structural data for the two RBD-targeting nAbs analyzed suggest comparable modes of recognition and neutralization for antibodies N-612-017 and N-612-056, which were selected from mRNA display, as those identified in convalescent or vaccinated donors.

### Structural Characterization of S1-Specific Antibodies N-612-014 and N-612-004

The primary target of SARS-CoV-2 nAbs is the viral spike glycoprotein, with the majority of nAbs targeting the RBD (Greaney et al., 2021a; Piccoli et al., 2020). The S NTD represents a common site of antigenic drift (Cele et al., 2021; McCarthy et al., 2021; Ribes et al., 2021), and nAbs that bind to this region have recently been identified (Cerutti et al., 2021; McCallum et al., 2021b; Suryadevara et al., 2021). To understand the binding mode of the NTD-targeting antibody N-612-014 (Figure 1), we determined a 3.5Å cryo-EM structure of N-612-014 Fabs complexed with stabilized S trimers (Figure 5A and Figure S2). N-612-014 adopted a binding pose parallel to the viral membrane and primarily used HC CDR loops to recognize an epitope at the periphery of the NTD (Figure 5A-C). The NTD epitope recognized by N-612-014 closely resembles that recognized by the human-derived SARS-CoV-2 antibody S2X316 that targets NTD antigenic site v, which resides outside of the antigenic supersite (site i) that is the main target of neutralizing NTD antibodies (McCallum et al., 2021b). The N-612-014 epitope (∼1070 Å^2^ epitope BSA) involves contacts with peripheral loops comprising NTD residues 68-78, 175- 188, and 245-260, as well as contacts with the tip of the supersite *β*-hairpin (Figure 5D and 5E). Despite contacts with residues 69-70NTD and 144NTD, N-612-014 maintains binding to S trimers of the B.1.1.7 lineage sequence (Figure S3), which has deletions at these positions that allow escape from NTD supersite antibodies (McCallum et al., 2021b). These data suggest that N-612- 014 retains NTD binding capability and may retain potency in the presence of NTD mutations commonly found in viral variants.

**Figure 5.**
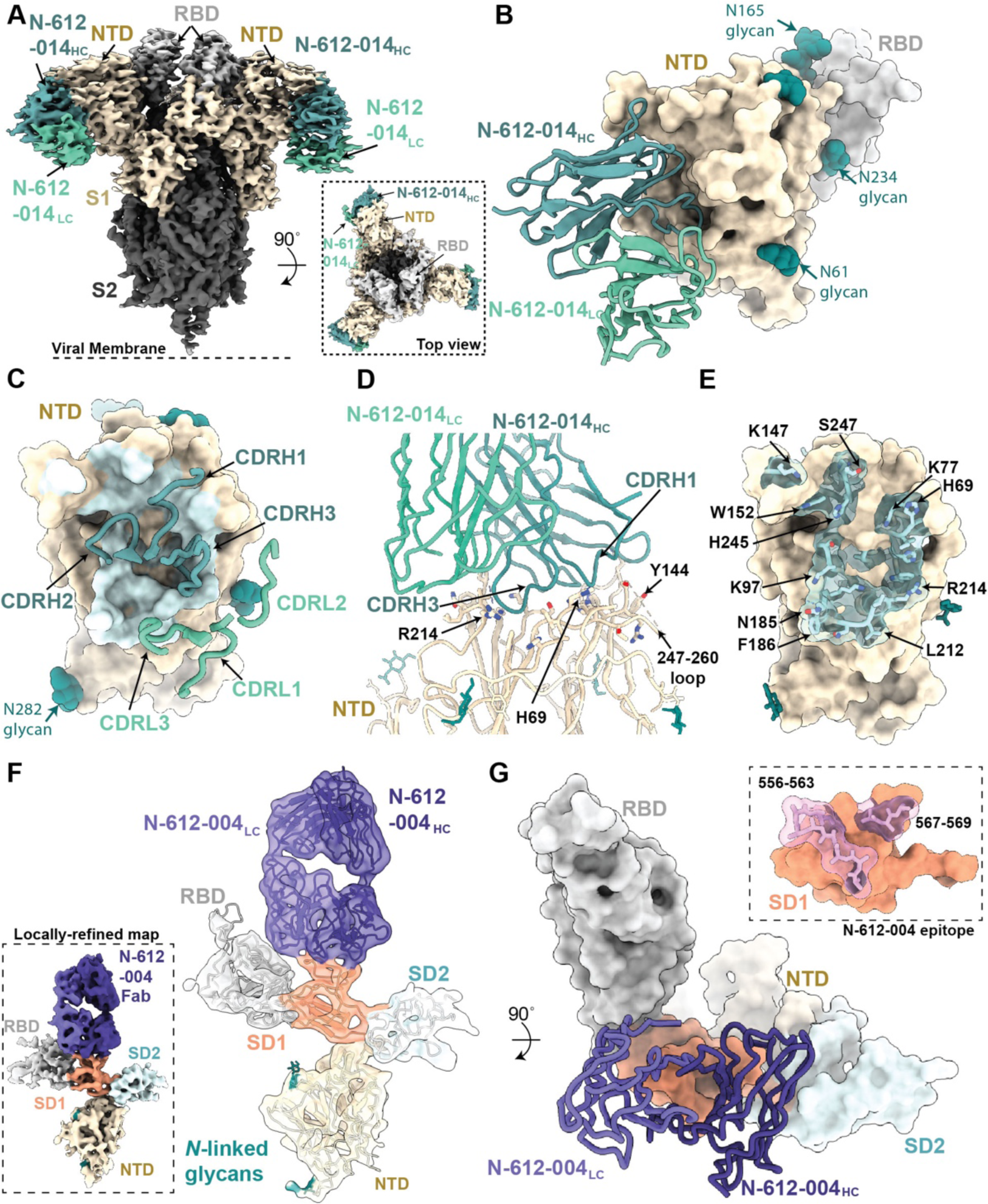
Structures of S1-Specific Antibodies N-612-014 and N-612-004 Bound to SARS-CoV-2 Spike. (A) Cryo-EM structure of the N-612-014 – S trimer complex. Inset: top down view of complex. (B) Close-up view of the N-612-014 variable domains (teal green) contacting the NTD (tan surface). The RBD (gray surface) of an adjacent protomer is shown as reference. (C) N-612-014 CDR loops (green ribbons) mapped onto the surface of the NTD (tan surface). Cartoon representation of the N-612-014 – NTD interface. (E) Surface and stick representation of the N-612-014 epitope (light green surface). (F) Cryo-EM structure of the N- 612-004 – S1 protomer (inset) rigid body fit with individual S1 domains (cartoon). (G) Cartoon and surface representation of N-612-004 (purple) recognition of the SD1 domain. Inset: N-612- 004 epitope (pink sticks) highlighted on the SD1 surface (orange). Given the low resolution, epitope residues were assigned using a criterion of a distance of ≤7 Å between antibody-antigen Cα atoms.

In addition to N-612-014, we also identified antibody N-612-004, an S1-specific antibody that was mapped to a domain outside of the NTD and RBD (Figure 1). Using single-particle cryo-EM, we determined a 4.8Å structure of N-612-004 bound to a dissociated S1 protomer, which revealed recognition of a SD1 epitope (Figure 5F and Figure S2). Consistent with our library design that varied CDR loops H2, H3 and L3, N-612-004 contacts were solely mediated by these three regions, which led to recognition of loops 556-563 and 567-69 in the SD1 domain (Figure 5G). The epitope for N-612-004 is not accessible on S trimers, which likely explains the lack of N-612-004-like antibodies identified among a repertoire of antibodies found in convalescent plasma (Figure S4).

### Activity of Identified nAbs Against Variants

To assess the relative affinity of RBD-binding nAbs N-612-017 and N-612-056 against a series of variants, BLI was performed using RBD variants B.1.1.7 (N501Y), B.1.351 (K417N/E484K/N501Y), CAL.20C (L452R), and A.VOI.V2(T478R/E484K) with single or combined mutations. N-612-056 binding affinity was not affected by any of the RBD mutations tested, which was anticipated based on structural characterization findings that indicate N-612- 056 recognizes a more conserved epitope on the surface of RBD. Neither N501Y (the only RBD mutation in B.1.1.7 and one of 3 RBD mutations in B.1.351) nor K417N, (one of 3 RBD mutations in B.1.351) disrupted the binding affinity of N-612-017. E484K - an escape mutant found in many different variants, including B.1.351 and A.VOI.V2 (Oliveira et al., 2021; Tegally et al., 2021; Weisblum et al., 2020) - did, however, reduce binding affinity of N-612-017 by 6-10 fold (Figure 6A and 6B). Furthermore, the L452R mutation found in CA.20C (also known as B.1.1.427 and B.1.1.429) completely abolished the binding of RBD by N-612-017.

**Figure 6.**
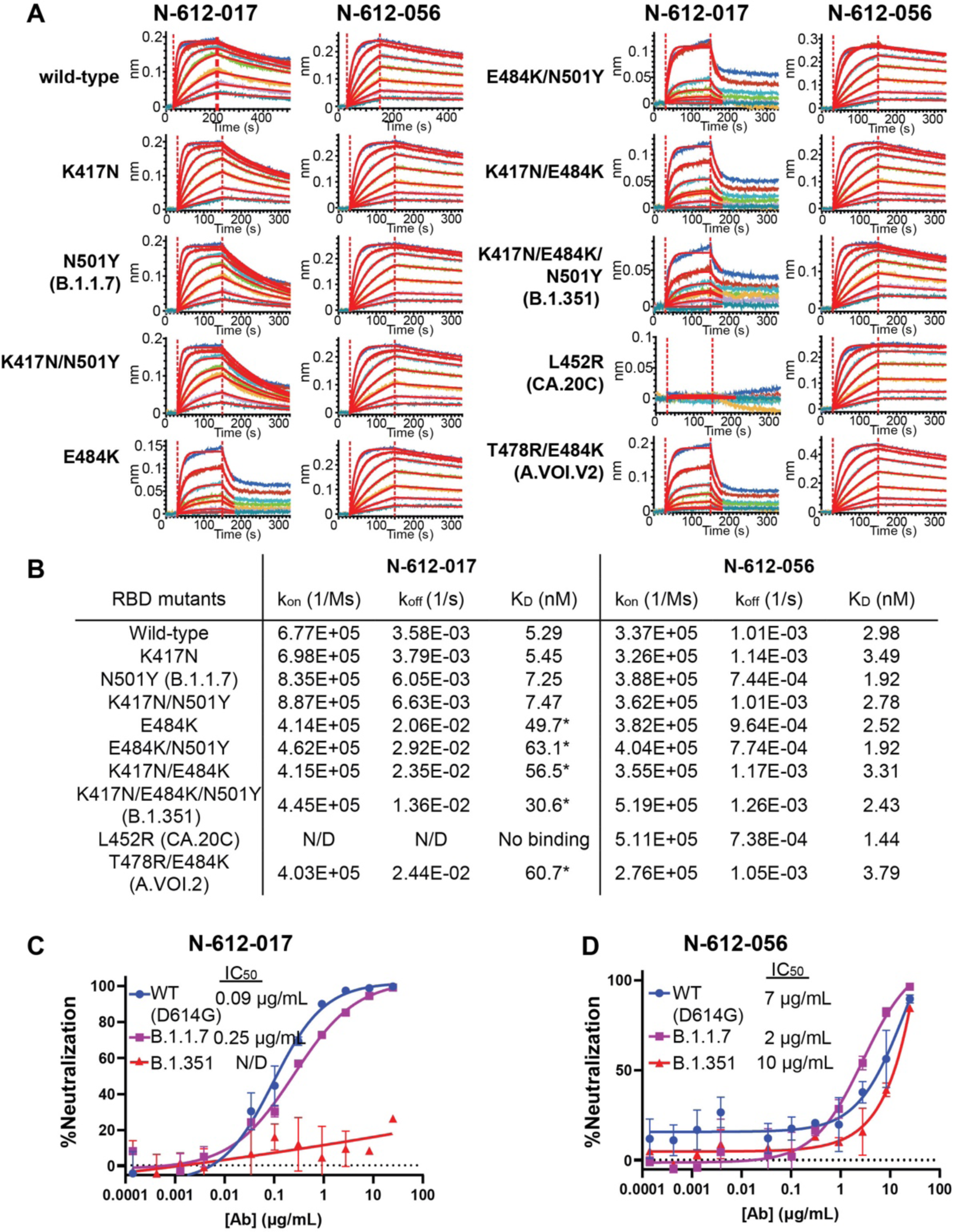
Binding Affinity and Neutralization Activity of N-612-017 and N-612-056 Against Known SARS-CoV-2 Variants. (A) BLI kinetic analysis of N-612-017 and N-612-056 affinity against various mutations found in SARS-CoV-2 variants alone or in combination. N-612-017 binding curves against RBD mutants containing E484K were fit with 1:1 binding model using a shorter dissociation time (30 sec) to highlight weakened binding. (B) Table of BLI kinetic assay values. SARS-CoV-2 pseudovirus neutralization assay of antibodies. Asterisk indicates *K*D values obtained from processing the data with a shorter dissociation time to fit the curves to 1:1 binding and may not represent accurate *K*D. (C) N-612-017 and (D) N-612-056 against wild-type (D614G), B.1.1.7, and B.1.351 variants. Mean and standard deviation of duplicate experiments (n=4), is shown.

N-612-017 and N-612-056 were then evaluated in a pseudovirus neutralization assay (Crawford et al., 2020) using wild-type (containing D614G), B.1.1.7, and B.1.351 pseudoviruses. N-612-017 neutralized wild-type (D614G) and B.1.1.7 pseudoviruses with IC50 = 0.09-0.25 µg/mL but failed to neutralize B.1.351. N-612-056 retained neutralization activity against all variants with IC50 of 2-10 µg/mL as expected (Figure 6C).

The binding affinity of N-612-014 and N-612-004 against the recombinant S1 domain containing B.1.1.7 mutations was tested and it was determined that 69-70del and Y144del on NTD did not affect binding affinity of N-612-014 for S1 whereas these mutations moderately lowered (by about 3-fold) the binding affinity of N-612-004 for S1 (Figure S3B and C).

### Generation of E484K and L452R-Resistant N-612-017

To recover N-612-017 binding against RBD with the E484K substitution (RBD-E484K), we used an mRNA-doped library for affinity maturation and identified mutations on VH framework 3 (Arg 71à Ser) and CDR H3 (Asp 97à Glu) that restored binding affinity against RBD- E484K. The N-612-017 subclone N-612-017-01 containing a single VH:R71S mutation and subclone N-612-017-03 containing double VH:R71S/D97E mutations were tested by BLI for their binding of RBD-B.1.351 and RBD-L452R. Interestingly, both N-612-017-01 and N-612- 017-03 subclones not only exhibited restored binding affinity against E484K-expressing RBD variants, they had 10- to 20-fold enhanced affinity against wild-type RBD (Figure 7A, 7B and Table S7). While neither subclone displayed complete recovery of binding affinity against RBD- L452R, affinity was relatively enhanced (*K*D = 33.1-58.7 nM). These subclones were then tested in a live virus neutralization assay against wild-type (D614G) and B.1.351 viruses and showed neutralization activity against both, whereas the parent N-612-017 did not show neutralization activity against B.1.351 (Figure 7C and 7D).

**Figure 7.**
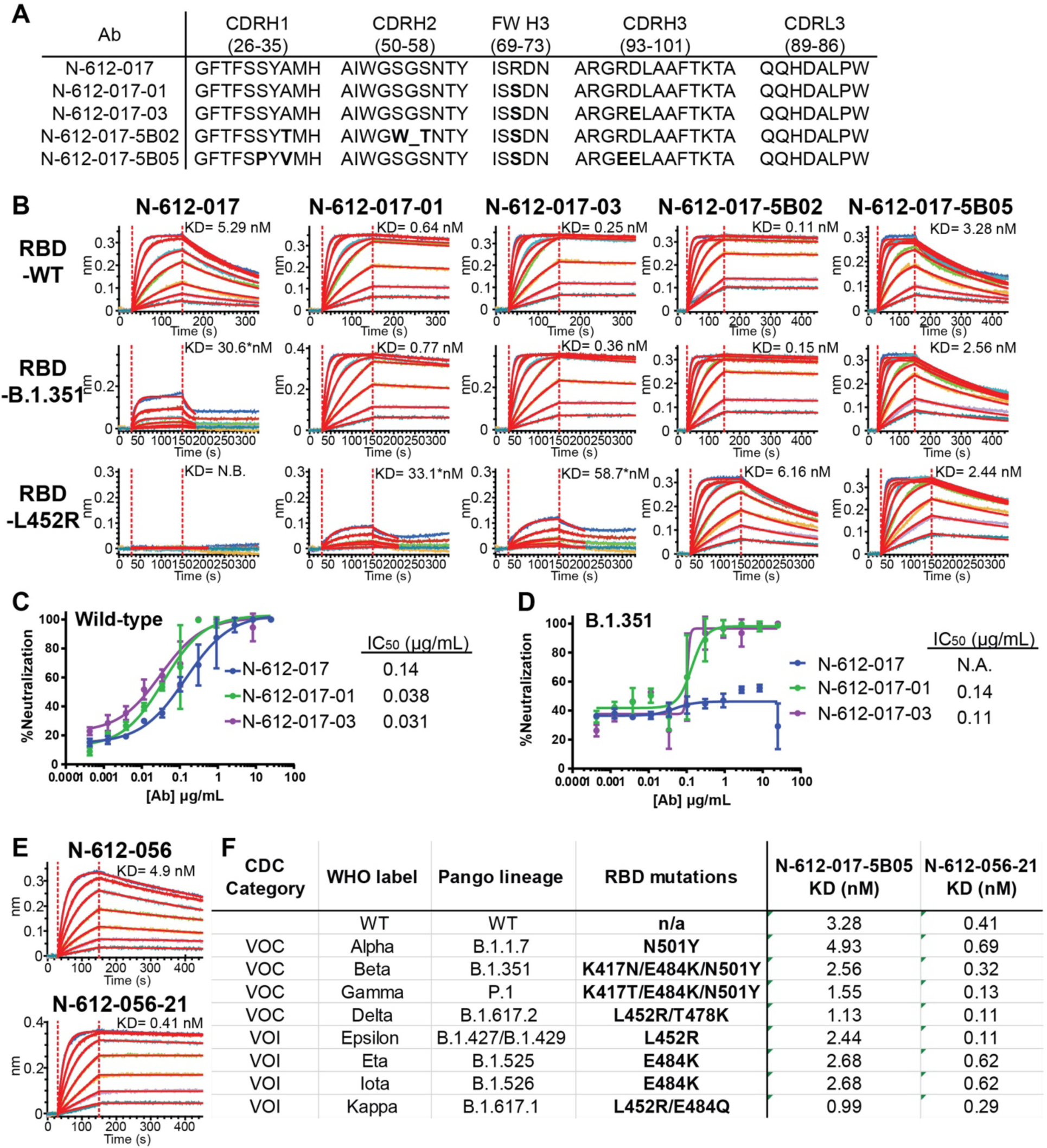
Affinity Maturation of N-612-017 and N-612-056. (A) VH and VL sequences of N- 612-017 affinity matured subclones. (B) BLI kinetic analysis of N-612-017 affinity matured subclones against RBD-wild-type, RBD-B.1.351, and RBD-L452R. Asterisk indicates *K*D values obtained from processing the data with a shorter dissociation time to fit the curves to 1:1 binding model and may not represent accurate *K*D values. SARS-CoV-2 live virus neutralization assay of N-612-017, N-617-017-01, and N-612-017-03 against (C) wild-type and (D) B.1.351 variant. Mean and standard deviation of duplicate experiments (n=3) (E) BLI kinetic analysis of N-612- 056 and affinity matured N-612-056-21 against RBD-wild-type. (F) Table of binding affinity of N-612-017-5B05 and N-612-056-21 against RBD containing variant mutations from all the VOC and VOI listed by CDC.

Subsequently, we used N-612-017-001 in affinity maturation against RBD-L452R and identified 2 clones with restored affinity against RBD-L452R. N-612-017-5B02 containing 4 additional VH mutations (A33T/S54W/G54Δ/S55T) and N-612-017-5B05 containing 4 additional VH mutations (S31P/A33V/R96E/D97E) were tested by BLI for their binding of B.1.351 and RBD-L452R. Both subclones showed complete recovery of binding affinity against RBD-B.1.351 and RBD-L452R (Figure 7A, 7B, and Table S7).

### Affinity Maturation of N-612-056

To improve potency of N-612-056, we utilized mRNA display for affinity maturation and identified N-612-056-21 containing a single point mutation in VH CDR3 (Ser 99 à Pro) that resulted in a 10-fold improvement in binding affinity (*K*D= 0.41 nM) (Figure 7E).

Affinity matured N-612-017-5B05 and N-612-056-21 were tested against all the variants of concern (VOC) and variants of interest (VOI) using BLI. N-612-017-5B05 showed binding affinity to the variants that was similar to the parent molecule N-612-017, whereas N-612-056-21 displayed binding affinities improved as much as ∼10-fold compared to N-612-056 (Figure 7F).

## Discussion

Our use of *in vitro* mRNA display facilitated our identification of novel antibody sequences and enabled us to enhance their binding affinity for mutated S of SARS-CoV-2 variants through affinity maturation. BLI and epitope binning analysis determined that 10 unique IgG1 antibody sequences identified here bind 7 distinct epitope regions on SARS-CoV-2 spike protein. Major sequence differences in CDRH3 and CDRL3 loops in combination with minor variation in CDRH1 and CDRH2 can drive the recognition of a broad spectrum of epitopes and create potent neutralizing interactions with SARS-CoV-2 spike protein. Previous IGHV (immunoglobulin heavy chain variable region) gene analysis identified distinct IGVH genes (e.g. 1-53) that were more likely to produce potent RBD-binding nAbs (Dejnirattisai et al., 2021; Robbiani et al., 2020; Yuan et al., 2020) within the human antibody repertoire. The nAb N-612-017 VH sequence is most similar to IGHV 3-23, which are most abundant in human antibodies, and shows potent neutralizing activity (< 0.1 µg/mL), suggesting CDR sequence variation is essential in determining potency against antigen regardless of germline genes.

The majority of potent nAbs (∼90%) are RBD targeting and the remainder target NTD of spike protein (Brouwer et al., 2020; Cao et al., 2020; Liu et al., 2020; McCallum et al., 2021b). Our most potent nAb N-612-017 is RBD targeting and categorized into Class 2 as characterized by cryo-EM. Though N-612-056 also neutralized SARS-CoV-2 by targeting RBD, albeit at lower potency, it lacks the ability to directly block ACE2 binding and binds to a cryptic epitope on RBD. Similar antibody binding to this cryptic epitope by a patient-derived antibody has been previously reported (Dejnirattisai et al., 2021). The rarity of this epitope is evident in the convalescent plasma blocking assay, in which convalescent plasma from 3 out 4 patients failed to block N-612-056 from binding to spike protein. This cryptic interface is well conserved, and although the potency of N-612-056 is relatively low, cross-reactivity with SARS-CoV RBD and sustained binding affinity for mutant SARS-CoV-2 RBD found in circulating variants suggests N-612-056 may be an attractive monoclonal antibody therapy candidate against novel variants.

The cryo-EM structural data presented here reveal that N-612-014 binds NTD at a site different from that of the majority of described antibodies (McCallum et al., 2021b), suggesting the presence of a second neutralization site on the NTD. Proposed mechanisms for nAbs targeting NTD include destabilization of S trimer by S1 shedding and blockage of cell-cell fusion auxiliary receptor binding, membrane fusion, or proteolytic activation (Huo et al., 2020; Walls et al., 2019; Wec et al., 2020; Wrobel et al., 2020). N-612-014 displayed neutralization activity in live virus assays whereas it lacked neutralization activity in a pseudovirus assay. Although this type of discrepancy is rare, such inter-assay discrepancies have been described previously (Liu et al., 2020). N-612-014 may require a longer incubation time to reach maximum neutralization because this allows the opportunity for the S trimer to adopt a conformation that is more susceptible to S1 shedding that is promoted by the antibody, thus destabilizing spike; this hypothesis awaits experimental confirmation.

The SD1-targeting antibody N-612-004 displayed partial neutralization activity and was only observed in complex with S1 domain dissociated from the spike trimer in cryo-EM. To our knowledge, there have been no reports on SD1-targeting antibodies that display neutralization activity. We also identified the S2-targeting antibody N-612-007 that displayed partial neutralization activity in a live virus neutralization assay and while structural analysis was attempted, we were unable to visualize/characterize an S trimer/N-612-007 complex. nAbs targeting S2 domain have been previously observed in MERS-CoV and SARS-CoV (Elshabrawy et al., 2012; Lai et al., 2005; Lip et al., 2006; Pallesen et al., 2017), and recently reported structures of SARS-CoV-2 S2/antibody complex have also revealed antibody binding to the S2 stem helix, which may interfere with membrane fusion machinary (Zhou et al., 2021). Other S2 epitope regions identified for SARS-CoV nAb are two heptad repeats region essential in cell fusion during virus entry (Elshabrawy et al., 2012; Lip et al., 2006; Pallesen et al., 2017).

Neutralization activities of these non-RBD binders (N-612-014, N-612-004, and N-612-007) were inconsistent between multiple assays and generally not very potent when tested individually. However, when tested in combination with N-612-017, all slightly enhanced the neutralization activity of N-612-017. This may suggest the role of non-RBD binding antibodies in neutralization.

Bamlanivimab is a Class 2 RBD binder that neutralizes wild-type SARS-CoV-2 and was the first antibody to attain emergency use authorization (EUA) by the FDA (Jones et al., 2021). This EUA was, however, recently revoked due to loss of potency against SARS-CoV-2 variants (Widera et al., 2021). Two alternate monoclonal antibody therapies remain available under EUA: a combination of casirivimab plus imdevimab (Baum et al., 2020; Pinto et al., 2020) and combination bamlanivimab plus etesevimab. In both cases, administering 2 monoclonal antibodies together is key to compounding potency and reducing the risk of variant virus escape from neutralization. Our N-612-017 antibody is a Class 2 RBD binder similar to bamlanivimab that also displays a loss of activity in the presence of the E484K mutation. To address this potential loss of efficacy, we used affinity maturation for N-612-017 and quickly identified subclones that restored affinity for both E484K and L452R. N-612-056 is resistant to all the RBD variants and N-612-014 is not affected by NTD mutation present in B.1.1.7. These nAbs are attractive candidates for the use in combination with N-612-017. N-612-056 was quickly affinity matured to present an attractive combo approach in combination with N-612-017 to combat variants.

The recent emergence of more transmissible and infectious variants such as B.1.351 (‘Beta’) and B.1.617 (‘Delta’) highlights the need for a method to rapidly address mutations that overcome current therapies and existing immunity. The results described in this study demonstrate the utility of mRNA display-based nAb discovery in the identification of antiviral monoclonal antibodies against the rapidly evolving SARS-CoV-2 pathogen which should be applicable to other novel or seasonal pathogens.

## Acknowledgements

We thank J. Vielmetter, P. Hoffman, and the Protein Expression Center in the Beckman Institute at Caltech for expression assistance and K. Huey-Tubman for assistance with soluble spike purification. Electron microscopy was performed in the Caltech Cryo-EM Center with assistance from S. Chen and A. Malyutin. We thank the Gordon and Betty Moore and Beckman Foundations for gifts to Caltech to support the Molecular Observatory. We thank J. Kaiser, director of the Molecular Observatory at Caltech, and beamline staff C. Smith and S. Russi at SSRL for data collection assistance. Use of the Stanford Synchrotron Radiation Lightsource, SLAC National Accelerator Laboratory, is supported by the U.S. Department of Energy, Office of Science, Office of Basic Energy Sciences under Contract No. DE-AC02-76SF00515. The SSRL Structural Molecular Biology Program is supported by the DOE Office of Biological and Environmental Research, and by the National Institutes of Health, National Institute of General Medical Sciences (P30GM133894). The contents of this publication are solely the responsibility of the authors and do not necessarily represent the official views of NIGMS or NIH. This work was supported by NIH (P01-AI138938-S1 to P.J.B.), the Caltech Merkin Institute for Translational Research (P.J.B.), and a George Mason University Fast Grant (P.J.B.). C.O.B was supported by the Hanna Gray Fellowship Program from the Howard Hughes Medical Institute and the Postdoctoral Enrichment Program from the Burroughs Wellcome Fund.

## Author contributions

Experimental designs: C.A.O., S.T, C.O.B, K.N., S.R., and P.S.S; mRNA library display: C.A.O; Cloning: C.A.O and W.H.; Protein Expression and Purification: C.A.O., S.T. and M.G.; Kinetic Analysis, ACE2 Blocking Assay, Epitope Binning, and ELISA ACE2 blocking assay: S.T., cryo-EM and X-ray crystallography: C.O.B and P.J.B.; Vero E6 Live Virus Neutralization Assay: J.T., A.R., M.M.F. and D.B.; pseudo-typed virus neutralization assay: P.G.. Manuscript preparation: S.T., C.O.B., C.A.O., and P.S..

## Materials Availability

All expression plasmids generated in this study for CoV proteins, CoV pseudoviruses, human Fabs and IgGs are available upon request through a MTA.

### Data availability

The atomic model generated for the N-612-056 Fab complexed with SARS-CoV-2 RBD have been deposited in the Protein Data Bank (PDB, http://www.rcsb.org/) under accession code 7S0B. The atomic models and cryo-EM maps generated for the N-612-017, N-612-014, and N-612-004 Fabs complexed with SARS-CoV-2 S have been deposited at the PDB (http://www.rcsb.org/) and the Electron Microscopy Databank (EMDB) (http://www.emdataresource.org/) under accession codes 7S0C, 7S0D, 7S0E and EMD-24786, EMD-24787, EMD-24788, respectively. All models and maps are publicly available as of the date of publication.

## Experimental Methods

### mRNA display

A synthetic VH3/Vk1 scFv library was transcribed by T7 run-off transcription (Thermo Fisher), followed by ligation to the pF30P linker (Liu et al., 2000) via a splint oligonucleotide by T4 DNA ligase (NEB). After lambda exonuclease digestion to remove splint and unincorporated linker, the ligated mRNA was purified by oligo(dT)25 dynabeads (Thermo Fisher). The mRNA-puromycin template was translated (Purexpress, NEB) followed by incubation with KCl (550 mM final) and MgCl2 (60 mM final) for 1 hour at room temperature to enhance fusion formation (Liu et al., 2000). The mRNA-scFv fusions were then affinity-purified using M2 anti-Flag beads (Sigma-Aldrich) to remove non-fused template and sequences containing nonsense mutations (Liao et al., 2009; Olson et al., 2011). After elution with 3XFlag peptide (Sigma-Aldrich), the fusions were reverse transcribed with super script II (ThermoFisher). The pool was incubated with biotinylated SARS CoV2 Spike extracellular domain bound to 5 μL streptavidin M280 dynabeads (ThermoFisher) for 1 hour at room temperature. After washing, the immobilized fusion samples were eluted by heat (95°C) and PCR amplified with KOD hot start polymerase (EMD). Affinity maturation was performed by replacing the wild type CDRs H1, H2, H3, and L3 with synthetic DNA cassettes derived from oligonucleotides doped at 6% (Hutchison et al., 1986) and performing 3-5 rounds of mRNA display as described above.

### Production antibodies and recombinant SARS-CoV-2 S domains

#### Molecular cloning

SARS-CoV-2 Spike ECD 1-1208 (682-GSAS-685; 986-PP-987) fused to the T4 fibritin trimerization domain with C-terminal Avi- and His-tag were synthesized with gene block (IDT) and cloned into pcDNA.3 vector. RBD-SD1, wild type RBD and mutant RBD domains were subcloned into pcDNA.3 vector with C-terminal His-tag.

VH and VL sequences of candidate sequences were cloned into a pcDNA.3 based vector with dual CMV promotor harboring IgG1 heavy chain and light chain backbone using NEBuilder Hifi DNA Assembly Master Mix (New England Biolabs).

#### FectoPRO® transient transfection of antibodies

For transient expression of antibodies by FectoPRO® transfection, CHO-S cells in suspension were cultured in CD-CHO media supplemented with 8 mM L-glutamine in shaker flasks at 37℃ with 125 rpm rotation and 8 % CO2. One day before transfection, CHO-S cells were seeded at a density of 1 x 10^6^ cells/mL in 45 mL culture flask. On the day of transfection, 75 µL of FectoPRO® transfection reagent (PolyPlus-transfection®) was mixed with 5 mL of 15 µg/mL pcDNA3 plasmid DNA harboring antibody encoding sequence in CD-CHO media and incubated for 10 min at room temperature. The DNA/transfection reagent mixture was added to 45 mL of CHO-S culture and incubated at 37℃ with 5% CO2 and 125 rpm rotation. On Day 3, 50 mL of the CD-CHO media supplemented with 8 mM L-glutamine was added and the culture incubated for an additional 4 days.

#### Lipofectamine® transient transfection of RBD constructs

For transient expression of RBD-SD1, RBD wild-type and RBD mutants, 293T cells were cultured and and incubated at at 37℃ with 5% CO2. Plasmid harboring RBD constructs were mixed with lipofectamin 2000 (Life Technology) with 1:1 (v:v) ratio and incubated for 20 min at room temperature. The mixture was then added to the culture and incubated for 3-4 days.

#### Maxcyte® transient transfection of SARS-CoV-2-S ECD trimer

For transient expression of SARS-CoV-2 S di-Pro ECD timer by Maxcyte® transfection, CHO-S cells were cultured in suspension in CD-CHO media supplemented with 8 mM L-glutamine in shaker flasks at 37℃ with 125 rpm rotation and 8% CO2. For transfection, cells in the exponential growth stage were pelleted by centrifugation at 1,400 rpm for 10 min, re-suspended in 10 mL of electroporation buffer, and re-pelleted at 1,400 rpm for 5 min. The cell pellet was resuspended at a density of 2 x 10^8^ cells/mL in electroporation buffer, mixed with the plasmid harboring SARS-CoV-2 S di-Pro ECD sequence at a concentration of 150 µg/mL, and transfected using OC-400 processing assemblies in a Maxcyte® ExPERT ATx Transfection System. Transfected cells were incubated for 30 min at 37℃, 5% CO2 and then resuspended in Efficient Feed A Cocktail (CHO-CD EfficientFeed™ A + 0.2% Pluronic F-68 + 1% HT Supplement + 1% L-glutamine) at a density of ∼4-6 x 10^6^ cells/mL. This cell culture was incubated at 37℃ with 5% CO2 and 125 rpm rotation overnight, 1 mM sodium buryrate was added, and the culture was further incubated at 32℃ with 3% CO2 and 125 rpm for 13 more days; during this incubation period, Maxcyte® Feed Cocktail (13.9% CD Hydrolysate, 69.5% CHO CD EfficientFeed™ A, 6.2% Glucose, 6.9% FunctionMax™ Titer Enhancer, 3.5% L- Glutamine) was added at 10% of the culture volume on Days 3 and Day 8.

#### Purification of IgGs

FectoPRO® transfection cell culture medium was centrifuged and filtered through a 0.22 µm filter to remove cells and debris, then loaded onto a HiTrap™ MabSelect SuRe™ column (GE Healthcare Life Sciences) on the AKTA Pure system pre-equilibrated with 10 mM Na Phosphate and 150 mM NaCl at pH 7.0. After loading, the column was washed with 10 column volumes of the same buffer. The protein was eluted with 100 mM sodium acetate, pH 3.6, then immediately neutralized using 2 M Tris pH 8.0. The elution fractions were pooled and dialyzed into 10 mM Hepes and 150 mM sodium chloride at pH 7.4.

#### Purification of Fabs

Fabs were generated by papain digestion using crystallized papain (Sigma-Aldrich) in 50 mMsodium phosphate, 2 mM EDTA, 10 mM L-cysteine, pH 7.4 for 30-60 min at 37℃ at a 1:100 enzyme:IgG ratio. Fab and partially cleaved IgG were applied on 1-mL HiTrap Protein L column (GE Healthcare Life Science). After loading, the column was washed with 10 column volumes of 10 mM Na Phosphate and 150 mM NaCl at pH 7.0. The protein was eluted with 100 mM sodium citrate, pH 2.5, then immediately neutralized using 2 M Tris pH 8.0. The elution fractions were pooled and dialyzed into 10 mM Hepes and 150 mM sodium chloride at pH 7.4. Fabs were further purified by SEC using a Superdex 200 10/300 GL column (GE Healthcare Life Sciences) in 10 mM Hepes and 150 mM sodium chloride at pH 7.4.

#### Purification of di-Pro S timer, RBD-SD1, RBD wild-type and RBD mutants

The Lipofectamin transfection cell culture medium and Maxcyte transfection cell culture medium was centrifuged and filtered through a 0.22 µm filter or 0.45 µm, respectively, remove cells and debris. 50 mM Tris, 100 mM sodium chloride, and 10 mM imidazole was added to the supernatant then loaded to a gravity column packed with Ni-NTA resins (Qiagen) pre-equilibrated with 20 mM Tris, 300 mM sodium chloride, and 10 mM imidazole, pH 8.0. After loading, the column was washed with 10 column volumes of the same buffer. The protein was eluted with 20 mM Tris, 150 mM sodium chloride, and 300 mM imidazole. The elution fractions were pooled and dialyzed into 10 mM Hepes and 150 mM sodium chloride, pH 7.4.

### Bio-Layer Interferometry (BLI) Kinetic Analysis of Antibodies

BLI buffer used in all experiments was 10 mM Hepes, 150 mM NaCl, pH 7.4, with 0.02% Tween 20, and 0.1% BSA. Analytes used in kinetic analysis were uncleavable S trimer, RBD- SD1, RBD wild-type, RBD mutants, commercially purchased recombinant SARS-CoV-2 S2 (SinoBiological), and SARS-CoV-2 S1 (SinoBiological). For determining binding affinities, IgGs were immobilized on Anti-hIgG Fc Capture (AHC) biosensors (Sartorius Corporation) and a concentration series of 200, 100, 50, 25, 12.5, 6.25, 3.125 nM was used to determine the equilibrium dissociation constants (*K*D values) for RBD-SD1, RBD wild-type, RBD mutants, and S1using 1:1 binding curve fit. For some RBD mutants that weakened binding and showed biphasic dissociation, only 30 seconds dissociation curves were used to fit 1:1 binding model. A concentration series of 20, 10, 5, 2.5, 1.25, 3.13, 1.56 nM was used to determine apparent *K*D for uncleavable S trimer using bivalent model on Octet HT software. For determining 1:1 binding affinity for S2, S2-His-tag was immobilized on Anti-Penta-His (HIS1K) biosensors (Sartorius Corporation), and a concentration series of S2 binding mAb Fab at 200, 100, 50, 25, 12.5, 6.25, 3.125 nM was used.

### ACE2 blocking assay

RBD-His-tag at 5 µg/mL was first loaded on Ni-NTA (NTA) biosensors (Sartorius Corporation) for 15 sec and subsequently blocked with 5 µg/mL mAbs or BLI assay buffer for 5 min. BLI signal from ACE2 binding were measured by incubating RBD-coated/mAb blocked biosensors in 25 nM ACE2-IgG1Fc for 3 min.

### Epitope binning

For epitope binning using S1 domain, biotinylated S1 binding mAbs at 25 µg/mL were first loaded on High Precision Streptavidin SAX biosensors (Sartorius Corporation) for 10 sec. 3.75 µg/mL of recombinant SARS-CoV-2 S1 were used to bind mAb captured on biosensors for 3 min and subsequently 10 µg/mL S1 binding mAb were incubated with biosensors to observe binding competition and signal was recorded for 3 min. For epitope binning using S2 domain, recombinant SARS-CoV-2 S2-His-tag at 10 µg/mL was loaded on Anti-Penta-HIS (HIS1K) biosensors (Sartorius Corporation) for 1 min. 10 µg/mL of S2 binding mAbs were sequentially incubated with biosensors for 3-5 min to observe binding competition.

### Vero E6 neutralization assay

All aspects of the assay utilizing virus were performed in a BSL3 containment facility according to the ISMMS Conventional Biocontainment Facility SOPs for SARS-CoV-2 cell culture studies. Vero E6 cells were seeded into 96-well plates at 20,000 cells/well and cultured overnight at 37°C. The next day, 3-fold serial dilutions of mAbs were prepared in DMEM containing 2% FBS, 1% NEAAs, and 1% Pen-Strep (vDMEM). SARS-CoV-2 virus stock was prepared in vDMEM at 10,000 TCID50/mL, mixed 1:1 (v:v) with the mAb dilutions, and incubated for 30 min or 24 hr at 37°C. Media was removed from the Vero E6 cells, mAb-virus complexes were added and incubated at 37°C for 48 hours before fixation with 4% PFA. Fixed cells were stained for SARS-CoV-2 nucleocapsid protein to measure infection. The percent neutralization was calculated as 100-((sample of interest-[average of “no virus”]/[average of “virus only”])*100).

### Pseudovirus neutralization assays

Pseudoviruses based on HIV lentiviral particles were prepared as described (Robbiani et al., 2020). Three-fold serially diluted mAbs were incubated with SARS-CoV-2 pseudovirus for 1 hour at 37°C. After incubation with 293TACE2 cells for 48 hours at 37°C, cells were washed twice with PBS, lysed with Luciferase Cell Culture Lysis 5x reagent (Promega), and NanoLuc Luciferase activity in lysates was measured using the Nano-Glo Luciferase Assay System (Promega). Relative luminescence units (RLUs) were normalized to values derived from cells infected with pseudovirus in the absence of mAbs. Half-maximal inhibitory concentrations (IC50 values) for mAbs were determined using 4-parameter nonlinear regression (Prism, GraphPad).

### Cryo-EM sample preparation

N-612-004, N-612-014 and N-612-017 Fab-S complexes were assembled by incubating purified SARS-CoV-2 S trimer at a 1.1:1 molar excess of purified Fab per S protomer at RT for 20 min. Complex was mixed with F-octylmaltoside solution (Anatrace) to a final concentration of 0.02% w/v and then 3 µL were immediately applied to a 300 mesh, 1.2/1.3 QuantiFoil grid (Electron Microscopy Sciences) that had been freshly glow discharged for 30s at 20 mA using a PELCO easiGLOW (Ted Pella). The grid was blotted for 3s with Whatman No. 1 filter paper at 22°C and 100% humidity then vitrified in 100% liquid ethane using a Mark IV Vitrobot (FEI) and stored under liquid nitrogen.

### Cryo-EM structure determination of N-612-004, N-612-014, and N-612-017 Fab in complex with S-6P

Single-particle cryo-EM data were collected for Fab-S trimer complexes as previously described (Barnes et al., 2020b) Briefly, movies were collected on a 200 kV Talos Arctica transmission electron microscope (Thermo Fisher) equipped with a Gatan K3 Summit direct electron detector operating in counting mode. Movies were collected using a 3x3 beam image shift pattern with SerialEM automated data collection software (Mastronarde, 2005) at a nominal magnification of 45,000x (super-resolution 0.4345 Å/pixel) using a defocus range of −0.7 to −2.0 µm. An average dose rate of 13.5 e^-^/pix/s resulted in a total dose of ∼60 e-/Å^2^ over 40 frames for all datasets.

For all datasets, movies were patch motion corrected for beam-induced motion including dose-weighting within cryoSPARC v3.1 (Punjani et al., 2017)after binning super resolution movies by 2 (0.869 Å/pixel). The non-dose-weighted images were used to estimate CTF parameters using Patch CTF in cryoSPARC, and micrographs with poor CTF fits, signs of crystalline ice, and field of views that were majority carbon were discarded. Particles were picked in a reference-free manner using Gaussian blob picker in cryoSPARC(Punjani et al., 2017) Initial particle stacks were extracted, binned x4 (3.48 Å/pixel), and subjected to *ab initio* volume generation (4 classes) and subsequent heterogeneous refinement with all particles. The 3D classes that showed features for a Fab-S trimer complex or Fab-S1 protomer were 2D classified to polish particle stacks. The resulting particle stacks were unbinned (0.869 Å/pixel) and re-extracted using a 432 box size, and moved to Relion v3.1 (Zivanov et al., 2018) for further 3D classification. Particles corresponding to distinct states were separately refined using non-uniform 3D refinement imposing C1 symmetry in cryoSPARC and final resolutions were estimated according to the gold-standard FSC (Bell et al., 2016).

To improve features at the Fab-RBD interface, focused, non-uniform 3D local refinement in cryoSPARC were performed by applying a soft mask around the Fab VHVL – RBD (N-612- 017), NTD (N-612-014), or SD1 (N-612-004) domains. These efforts resulted in a modest improvement in the Fab-S interface, which helped accurate model building.

### X-ray crystallography structure determination of N-612-056 in complex with RBD

The N-612-056-RBD complex was assembled by incubating the SARS-CoV-2 RBD with a 2x molar excess of Fab for 1 h on ice followed by size exclusion chromatography on an superdex200 10/300 increase column (Cytiva). Fractions containing complex were pooled and concentrated to 5-8 mg/mL. Crystallization trials using commercially-available screens (Hampton Research) were performed at room temperature using the sitting drop vapor diffusion method by mixing equal volumes of the Fab-RBD complex and reservoir using a TTP LabTech Mosquito instrument. Crystals were obtained for N-612-056-RBD complex in 0.2 M Lithium citrate tribasic tetrahydrate and 20% w/v polyethylene glycol 3,350, subsequently cryoprotected by adding glycerol directly to drops to a final concentration of 20% v/v and cryopreserved in liquid nitrogen.

X-ray diffraction data were collected at the Stanford Synchrotron Radiation Lightsource (SSRL) beamline 12-2 on a Pilatus 6M pixel detector (Dectris). Data from single crystals were indexed and integrated in XDS (Kabsch, 2010) and merged using AIMLESS in *CCP4* (Winn et al., 2011)(Table S6). The N-612-056-RBD structure was solved by molecular replacement in PHASER (McCoy et al., 2007) using unmodified RBD coordinates (PDB 7K8M) and coordinates from C002 Fab (PDB 7K8O) as search models, after removal of C002 heavy chain and light chain CDR loops. Coordinates were rigid body and B-factor refined in PHENIX v1.19 (Adams et al., 2010) followed by sequence matching and repeated cycles of *phenix.refine* and manual building in Coot (v0.9.3) (Emsley et al., 2010) (Table S6).

### Structure analyses

Buried surface area estimates were made using PDBePISA with a 1.4Å probe (Krissinel and Henrick, 2007). Potential hydrogen bonds were assigned using a distance of <3.6A and an A-D-H angle of >90 and the maximum distance allowed for a van der Waals interaction was 4.0 Å. Structure figures were made using UCSF Chimera v1.14 (Goddard et al., 2018).

## Supplementary Methods

### Developability assays

#### Meso Scale Diagnostics (MSD) Polyreactivity

Six different antigens, cardiolipin (50 µg/mL, C0563; Sigma), KLH (5 µg/mL, H8283; Sigma), LPS (10 µg/mL, tlrl-eblps; InvivoGen), ssDNA (1 µg/mL, D8899; Sigma), dsDNA (1 µg/mL, D4522; Sigma), and insulin (5 µg/mL, I9278; Sigma) were coated onto MSD MULTI-Array 96-well plate (MSD) individually at 50 µL per well overnight at 4°C. Plates were blocked with PBS with 0.5% BSA at room temperature for 1 h, followed by three washes with PBST (PBS plus 0.05% Tween 20). Fifty microliters of 100 nM testing antibody solution was added to each well and incubated at RT for 1 hour followed by six washes with 100 µL of PBS. Twenty microliters of 0.25 µg/mL SULFO-tag, anti-Human antibody was added to the wells and incubated for 1 hour followed by six washes as before. Finally, 150 µL of 2X MSD Read Buffer T (MSD) was added to each well, and electrochemilluminescence signal was read by MSD Sector Imager. Polyreactivity score was determined by normalizing signal by control wells with no test antibody.

#### Hydrophobic Interaction Column (HIC)

5 µg antibody samples (1 mg/mL) were spiked in with a mobile phase A solution (1.8 M ammonium sulfate and 0.1 M sodium phosphate at pH 6.5) to achieve a final ammonium sulfate concentration of about 1 M before analysis. A Sepax Proteomix HIC butyl-NP5 column on Agilent 1100 HPLC was used with a liner gradient of mobile phase A and mobile phase B solution (0.1 M sodium phosphate, pH 6.5) over 20 min at a flow rate of 1 mL/min with UV absorbance monitoring at 280 nm. Elution time was recorded.

#### Clone Self-interaction by Bio-layer Interferometry (CSI-BLI)

Human IgG (Sigma) was loaded to an AHQ biosensor (ForteBio) to ∼1 nm, followed by sensor blocking with human IgG1 Fc (R&D systems). The self-association was performed at 1 µM solution concentration of antibodies for 300s on an Octet Red96e system (Sartorius Corporation). The binding response from the association step was subtracted from that of a reference IgG.

*Accelerated Stability Assay*: Antibody samples at 1 mg/mL were kept at 40°C for 30 days in 10 mM Hepes and 150 mM sodium chloride, pH 7.4. 10 µg of antibody was loaded onto Zenix-C SEC-300 size-exclusion column (Sepax) on HPLC at Day 0, 5, 20, and 30. A long-term stability slope (% aggregation/day) was calculated from the percent aggregated measured on the SEC-HPLC at each time-point.

#### Differential Scanning Fluorimetry (DSF) Analysis of Melting Temperature (Tm)

Twenty microliters of 1mg/mL antibody sample was mixed with 10 µL of 20x SYPRO Orange (ThermoFisher) in a 96-well PCR plate (ThermoFisher). The plate was scanned from 40℃ to 95℃ at a rate of 0.5℃/ 2 min in a CFX96 Real-Time PCR system (Bio-Rad). The Fab Tm was assigned using the first derivative of the raw data.

### Convalescent plasma blocking assay

Spike trimer with C-terminal biotin at 5 µg/mL was first loaded on High Precision Streptavidin SAX biosensors (Sartorius Corporation) for 75 min. Spike coated biosensor was subsequently blocked with 10-fold diluted SARS-CoV-2 convalescent plasma for 15 min. BLI signal from 10 mAbs binding to available surface of spike timer were measure by incubating Spike-coated/plasma blocked biosensors in 10 µg/mL of mAbs for 3 min. BLI signal was compared to self-blocking of N-612-017/N-612-017 and non-blocking pair of N-612-017/N-612- 004 to determine whether each mAb was completely blocked, partially blocked, or non-blocked by convalescent plasma from 4 different patients.

## Supplementary Tables

**Table S1.**
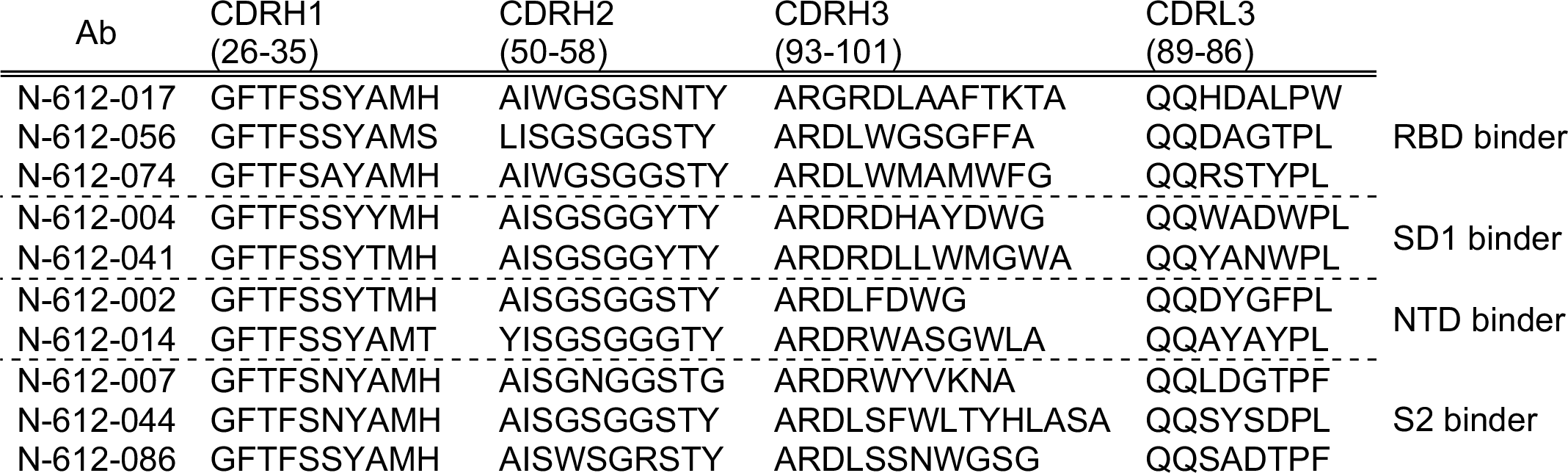
Ten unique spike binding VH/VL sequences identified by mRNA display

**Table S2.** BLI kinetic parameters obtained for various SARS-CoV-2 Spike domains (RBD, RBD-SD1, S1, S2)

| Analyte | nAb | $k_{on}$ (1/Ms) | $k_{off}$ (1/s) | $K_D$ (nM) |
| --- | --- | --- | --- | --- |
| RBD | N-612-017 | 6.77E+05 | 3.58E-03 | 5.29 |
|  | N-612-056 | 3.37E+05 | 1.01E-03 | 2.98 |
|  | N-612-074 | 1.33E+05 | 1.48E-03 | 11.1 |
| RBD-SD1 | N-612-017 | 4.56E+05 | 3.94E-03 | 8.65 |
|  | N-612-056 | 3.59E+05 | 1.04E-03 | 2.89 |
|  | N-612-074 | 1.21E+05 | 3.97E-03 | 32.9 |
|  | N-612-004 | 1.51E+05 | 9.79E-04 | 6.49 |
|  | N-612-041 | 2.10E+05 | 3.35E-03 | 16.0 |
| S1 | N-612-017 | 2.77E+05 | 2.04E-03 | 7.37 |
|  | N-612-056 | 1.51E+05 | 5.86E-04 | 3.89 |
|  | N-612-074 | 1.01E+05 | 1.46E-03 | 14.4 |
|  | N-612-004 | 9.92E+04 | 2.13E-04 | 2.15 |
|  | N-612-041 | 1.25E+05 | 1.83E-03 | 14.4 |
|  | N-612-002 | 2.68E+05 | 7.26E-04 | 2.71 |
|  | N-612-014 | 1.76E+05 | 2.18E-03 | 12.4 |
| S2 (Ligand) | N-612-007 (Fab) | 3.70E+05 | 1.40E-03 | 3.79 |
|  | N-612-044 (Fab) | 1.70E+05 | 1.61E-03 | 9.44 |
|  | N-612-086 (Fab) | 2.67E+05 | 0.96E-02 | 11.6 |

**Table S3.** BLI kinetic parameters obtained for Spike trimer using bivalent model fit

| **Ab** | **$k_{on1}$ (1/Ms)** | **$k_{on2}$ (1/Ms)** | **$k_{off1}$ (1/s)** | **$k_{off2}$ (1/s)** | **Apparent $K_D$ (nM)** |
| --- | --- | --- | --- | --- | --- |
| N-612-017 | 8.35E+04 | 1.55E+00 | <1.0E-07 | 1.69E-02 | <0.001 |
| N-612-056 | 6.24E+04 | 1.94E+00 | <1.0E-07 | 5.15E-02 | <0.001 |
| N-612-074 | 9.23E+04 | 8.12E-01 | <1.0E-07 | 1.89E-01 | <0.001 |
| N-612-004 | 7.60E+04 | 4.24E+00 | <1.0E-07 | 1.00E+00 | <0.001 |
| N-612-041 | 1.11E+05 | 4.06E-01 | <1.0E-07 | 1.75E-01 | <0.001 |
| N-612-002 | 5.69E+04 | 6.82E-02 | <1.0E-07 | 6.76E-02 | <0.001 |
| N-612-014 | 2.54E+04 | 3.82E-03 | 2.29E-07 | 2.82E+00 | 0.0033 |
| N-612-007 | 5.78E+04 | 7.80E-02 | <1.0E-07 | 2.92E-02 | <0.001 |
| N-612-044 | 7.15E+04 | 2.62E+00 | <1.0E-07 | 7.14E-02 | <0.001 |
| N-612-086 | 6.11E+04 | 1.55E-01 | <1.0E-07 | 5.53E-02 | <0.001 |

**Table S4.** Developability assay summary table

| **Ab** | **Polyreactivity MSD (Fold-over-PBS)** | **HIC (min)** | **BLI-CSI (nm)** | **Accelerated stability % monomer increase/day** | **Fab Tm (°C)** |
| --- | --- | --- | --- | --- | --- |
| N-612-017 | 3 | 10.1 | -0.06 | 0.09 | 85 |
| N-612-056 | 7 | 11.5 | 0.00 | 0.10 | 81 |
| N-612-074 | 31 | **22.1** | -0.01 | 0.10 | 76 |
| N-612-004 | 14 | 14.9 | 0.05 | 0.13 | 85 |
| N-612-041 | 10 | **20.4** | 0.05 | **0.37** | 88 |
| N-612-002 | 22 | 13.9 | 0.08 | 0.18 | 82 |
| N-612-014 | 4 | 12.3 | 0.09 | 0.16 | 88 |
| N-612-007 | 10 | 11.8 | 0.04 | 0.11 | 82 |
| N-612-044 | 8 | **13.4/15.4** | -0.06 | 0.10 | 88/91 |
| N-612-086 | 10 | 13.2 | 0.07 | 0.11 | 88 |
| Acceptance criteria | <50 | <16 | <0.2 | <0.2 | >65 °C |

**Table S5.** Cryo-EM data collection and refinement statistics (related to Figures 3 and 5).

|  | N-612-017 Fab<br>SARS-CoV-2 S 6P | N-612-014 Fab<br>SARS-CoV-2 S 6P | N-612-004 Fab<br>SARS-CoV-2 S 6P |
| --- | --- | --- | --- |
| **PDB** | **7S0C** | **7S0D** | **7S0E** |
| **EMD** | **24786** | **24787** | **24788** |
| **Data collection conditions** |  |  |  |
| Microscope | Talos Arctica | Talos Arctica | Talos Arctica |
| Camera | Gatan K3 Summit | Gatan K3 Summit | Gatan K3 Summit |
| Magnification | 45,000x | 45,000x | 45,000x |
| Voltage (kV) | 200 | 200 | 200 |
| Recording mode | counting | counting | counting |
| Dose rate (e<sup>-</sup>/pixel/s) | 13.5 | 13.3 | 13.8 |
| Electron dose (e<sup>-</sup>/Å<sup>2</sup>) | 60 | 60 | 60 |
| Defocus range (µm) | 0.7 – 2.0 | 0.7 – 2.0 | 0.7 – 2.0 |
| Pixel size (Å) | 0.8689 | 0.8689 | 0.8689 |
| Micrographs collected | 2,585 | 3,791 | 3,717 |
| Micrographs used | 2,132 | 3,211 | 2,047 |
| Total extracted particles | 282,890 | 505,695 | 595,163 |
| Refined particles | 175,986 | 389,223 | 115,068 |
| Particles in final refinement | 108,746 | 137,684 | 107,271 |
| Symmetry imposed | C1 | C1 | C1 |
| FSC 0.143 (unmasked/masked) |  |  |  |
| unmasked | 4.4 Å | 6.4 Å | 7.1 Å |
| masked | 3.2 Å | 3.5 Å | 4.8 Å |
| **Refinement and Validation** |  |  |  |
| Initial model used | 6XKL | 6XKL | 6XKL |
| Number of atoms |  |  |  |
| Protein | 29,164 | 33,650 | 5,307 |
| Ligand | 434 | 873 | 28 |
| MapCC (global/local) | 0.81/0.78 | 0.79/0.76 | 0.87/0.69 |
| Map sharpening B-factor | 65.8 | 75.3 | 155 |
| R.m.s. deviations |  |  |  |
| Bond lengths (Å) | 0.01 | 0.005 | 0.02 |
| Bond angles (°) | 0.9 | 0.89 | 1.5 |
| MolProbity score | 2 | 2.2 | 2.46 |
| Clashscore (all atom) | 13.4 | 17.7 | 7.9 |
| Poor rotamers (%) | 0.2 | 0.1 | 0 |
| Ramachandran plot |  |  |  |
| Favored (%) | 94.7 | 93.4 | 92.8 |
| Allowed (%) | 5.1 | 6 | 6.3 |
| Disallowed (%) | 0.2 | 0.6 | 0.9 |

**Table S6.**
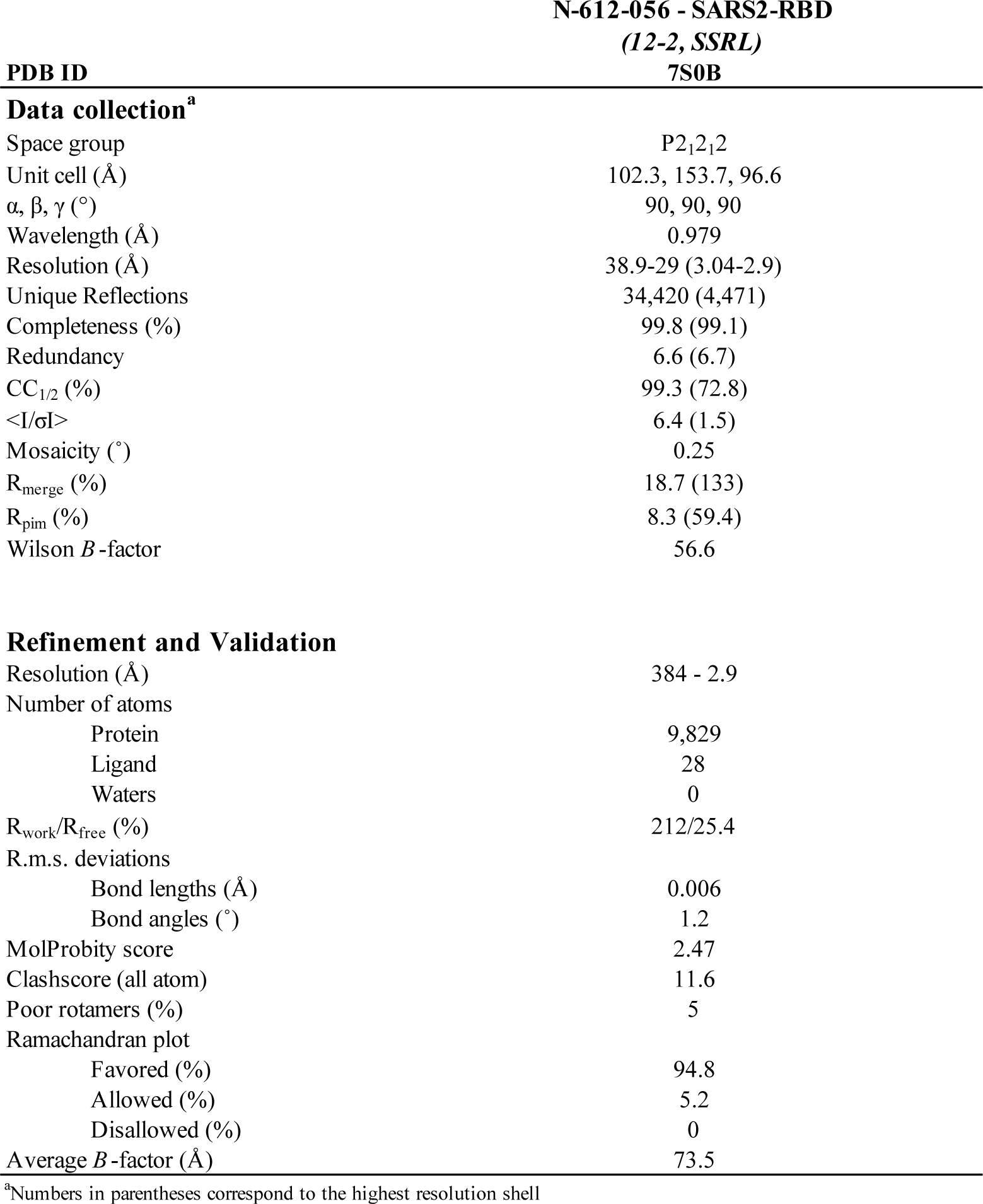
X-ray crystallography data collection and refinement statistics (related to **Figure 4**).

**Table S7.** BLI kinetic analysis of N-612-017, N-612-017-01, N-612-017-03, N-612-017-5B02, and N-612-017-5B05 against RBD-WT, RBD-B.1.351 (K417N/E484K/N501Y) and RBD- L452R

|  | RBD-WT |  |  | RBD-B.1.351 |  |  | RBD-L452R |  |  |
| --- | --- | --- | --- | --- | --- | --- | --- | --- | --- |
| | $k_{on}$ (1/Ms) | $k_{off}$ (1/s) | $K_D$ (nM) | $k_{on}$ (1/Ms) | $k_{off}$ (1/s) | $K_D$ (nM) | $k_{on}$ (1/Ms) | $k_{off}$ (1/s) | $K_D$ (nM) |
| N-612-017 | 6.77E+05 | 3.58E-03 | 5.29 | 4.45E+05 | 1.36E-02 | 30.6 | -- | -- | N.B. |
| N-612-017-01 | 5.59E+05 | 3.59E-04 | 0.64 | 5.31E+05 | 4.06E-04 | 0.77 | 2.34E+05 | 7.75E-03 | 33.1 |
| N-612-017-03 | 6.48E+05 | 1.60E-04 | 0.25 | 6.47E+05 | 2.33E-04 | 0.36 | 1.18E+05 | 6.90E-03 | 58.7 |
| N-612-017-5B02 | 8.24E+05 | 9.26E-05 | 0.11 | 7.99E+05 | 1.20E-04 | 0.15 | 5.74E+05 | 3.54E-03 | 6.16 |
| N-612-017-5B05 | 9.62E+05 | 3.16E-03 | 3.28 | 9.68E+05 | 2.48E-03 | 2.56 | 8.01E+05 | 1.96E-03 | 2.44 |

---

**Figure S1.**
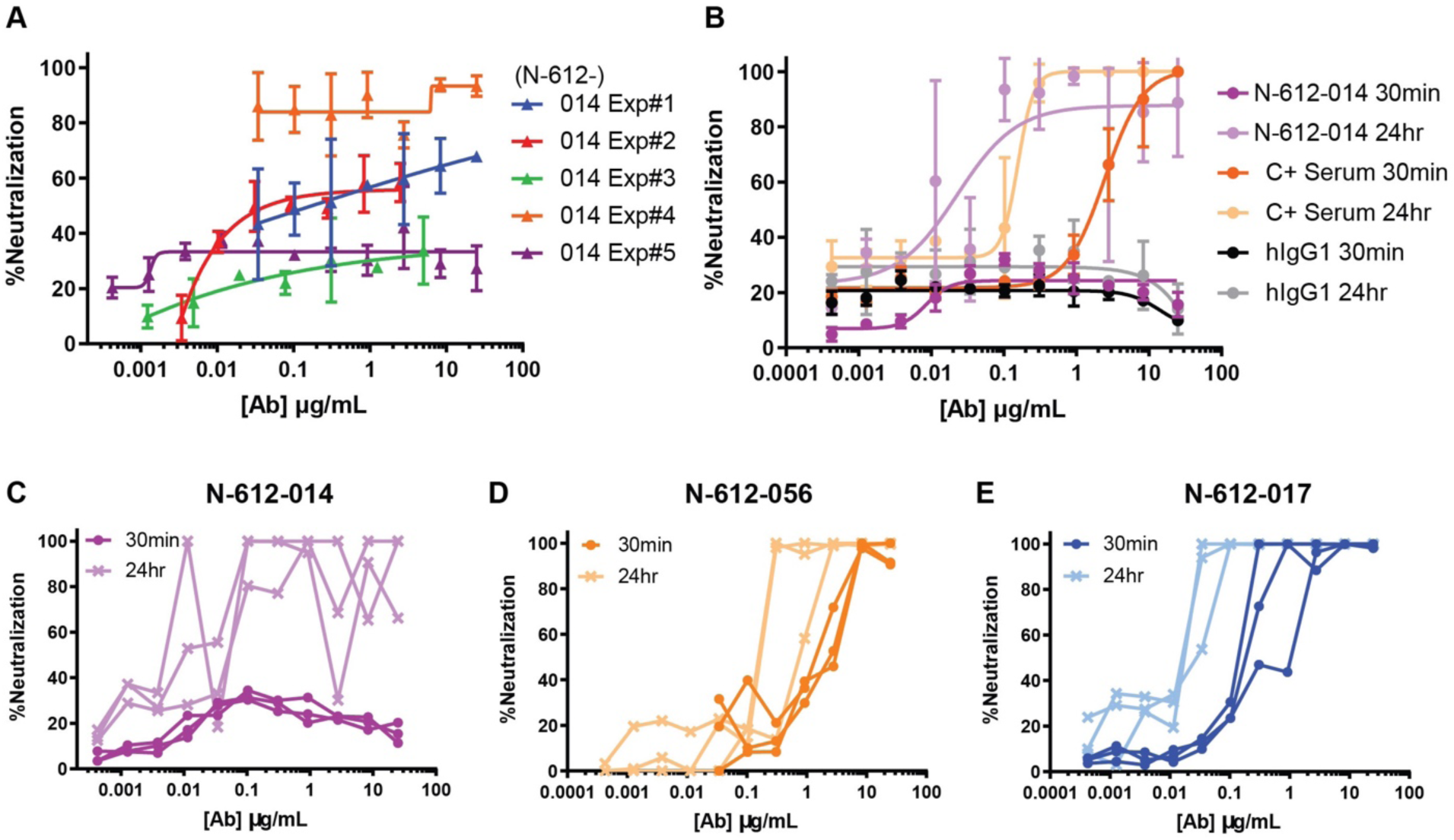
Vero E6 live virus neutralization assay. (A) Neutralization activity of N-612-014 in 5 separate experiments. (B) Comparison of neutralization activity of N-612-014, convalescent (C+) serum, and control human IgG with 30-min vs 24-hr antibody-virus incubation. (C, D, E) Change in neutralization potency by incubation time.

**Figure S2.**
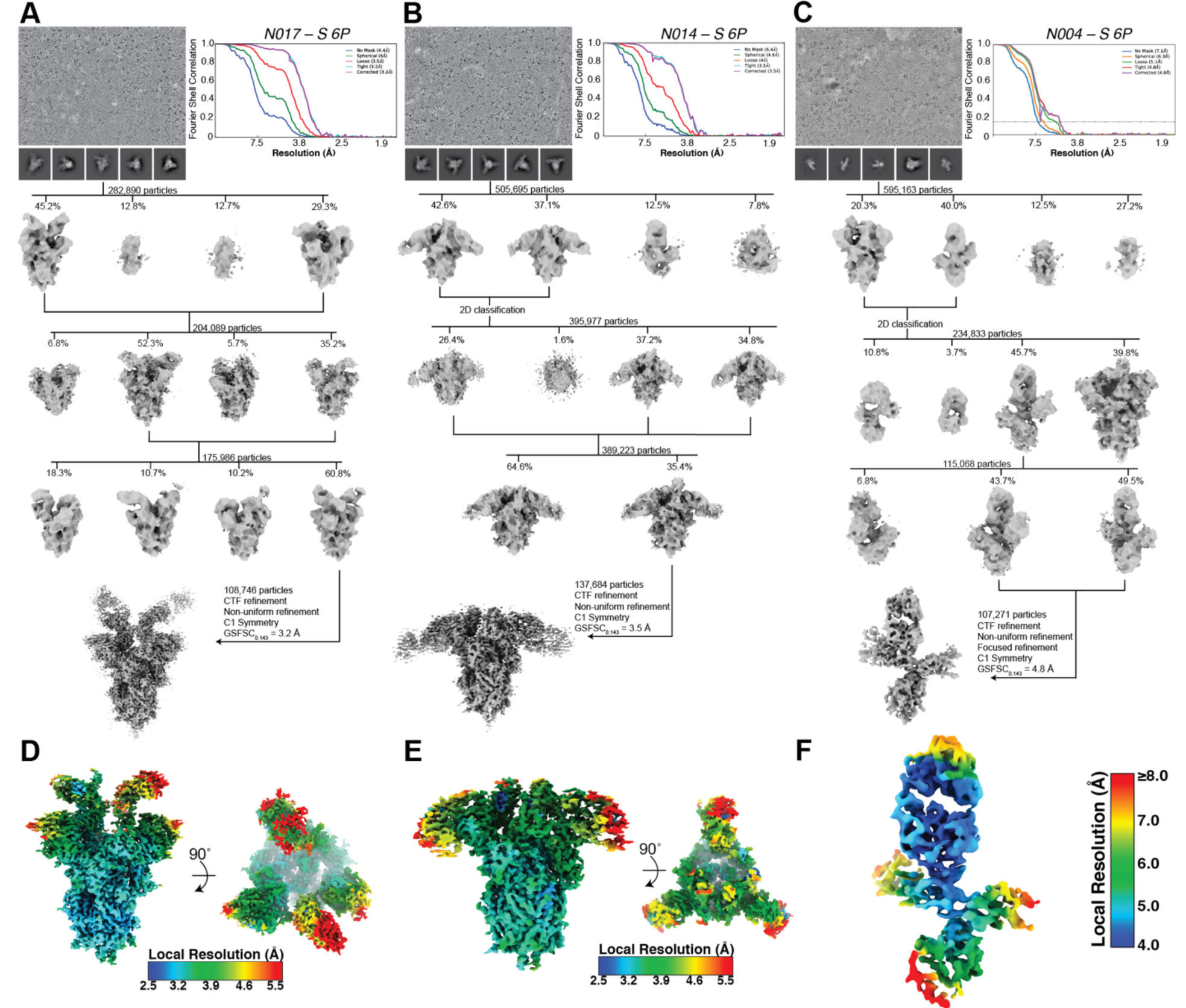
Cryo-EM data processing and validation (related to Figures 3 and 5). (A-C) Representative micrograph, 2D class averages, data processing workflow, and Gold Standard FSC plots for the final reconstructions of (A) N-612-017 – S 6P, (B) N-612-014 – S 6P, and (C) N-612-056 – S 6P complexes. (D-F) Local resolution estimates calculated in cryoSPARC v3.1 for (D) N-612-017 – S 6P, (E) N-612-014 – S 6P, and (F) N-612-056 – S 6P complexes.

**Figure S3:**
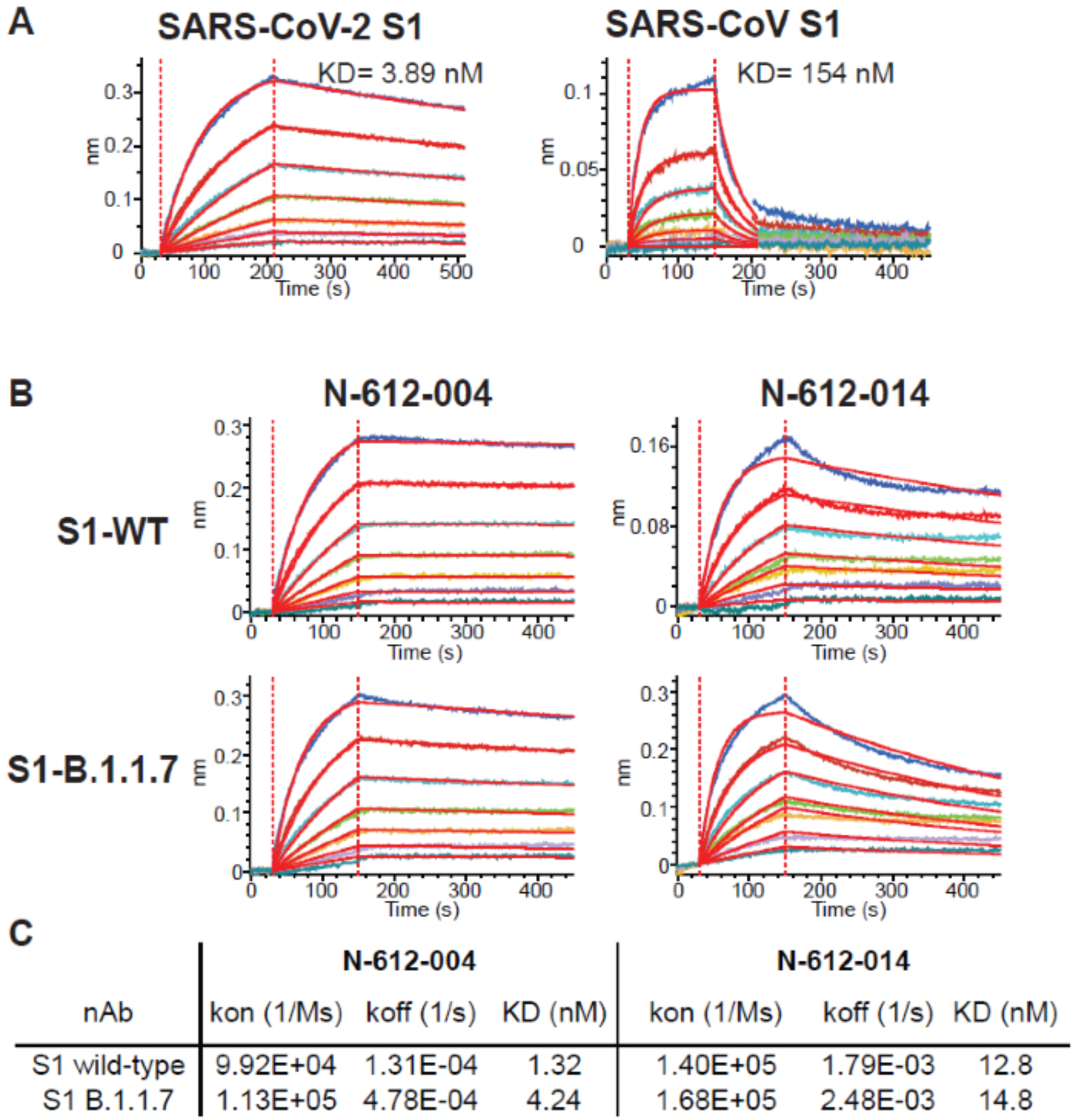
(A) Cross-reactivity of N-612-056 against SARS-CoV. BLI kinetic analysis of SARS-CoV-2 and SARS-CoV S1 domain binding to N-612-056. (B, C) BLI kinetic analysis of N-612-004 and N-612-014 against S1 domain from WT and B.1.1.7 variant of SARS-CoV-2 Spike protein.

**Figure S4.**
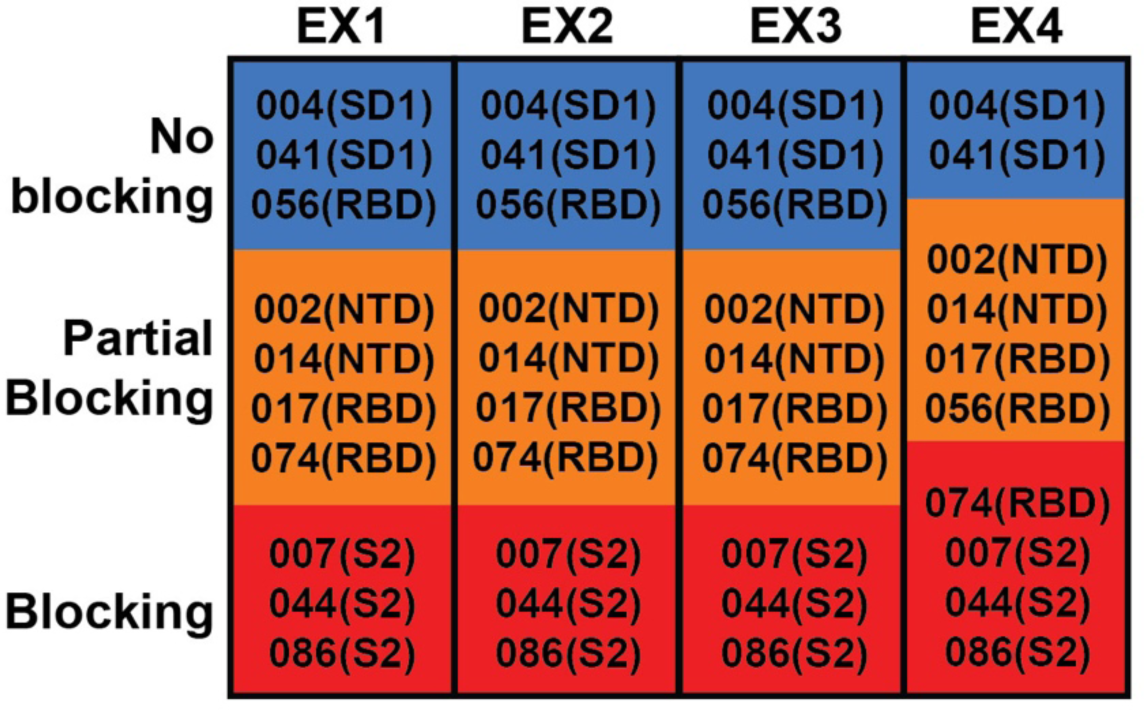
Convalescent plasma blocking of 10 mAbs binding to SARS-CoV-2 Spike.

